# An inexpensive, high-precision, modular spherical treadmill setup optimized for *Drosophila* experiments

**DOI:** 10.1101/2021.04.29.442008

**Authors:** Frank Loesche, Michael B. Reiser

## Abstract

To pursue a more mechanistic understanding of the neural control of behavior, many neuroethologists study animal behavior in controlled laboratory environments. One popular approach is to measure the movements of restrained animals while presenting controlled sensory stimulation. This approach is especially powerful when applied to genetic model organisms, such as *Drosophila melanogaster,* where modern genetic tools enable unprecedented access to the nervous system for activity monitoring or targeted manipulation. While there is a long history of measuring the behavior of body- and head-fixed insects walking on an air-supported ball, the methods typically require complex setups with many custom components. Here we present a compact, simplified setup for these experiments that achieves high-performance at low cost.

The simplified setup combines existing hardware and software solutions with new component designs. We replaced expensive optomechanical and custom machined components with off-the-shelf and 3D-printed parts. We built the system around an inexpensive camera that achieves 180 Hz imaging and use an inexpensive tablet computer for presenting view-angle-corrected stimuli updated through a local network. We quantify the performance of the integrated system and characterize the visually guided behavior of flies in response to a range of visual stimuli. The improved system is thoroughly documented. This publication is accompanied by CAD files, parts lists, source code, and step-by-step instructions for setting up the system and analyzing behavioral data. We detail a complete ~$.300 system, including a cold-anesthesia tethering stage, that is ideal for hands-on teaching laboratories. This represents a nearly 50-fold cost reduction as compared to a typical system used in research laboratories, yet is fully featured and yields excellent performance.

We report the current state of this system, which started with a one-day teaching lab for which we built seven parallel setups and continues towards a setup in our lab for larger-scale analysis of visual-motor behavior in flies. Because of the simplicity, compactness, and low cost of this system, we believe that high-performance measurements of tethered insect behavior should now be widely accessible and suitable for integration into many systems. This access enables broad opportunities for comparative work across labs, species, and behavioral paradigms.

## 1 INTRODUCTION

The fly *Drosophila melanogaster* is a powerful model system for research in nearly all areas of organismal biology, and has been especially central to major discoveries in the development and function of the nervous system (Bellen et al., 2010). *Drosophila* have long been champion species for a wide range of behavioral experiments that are ideally suited to a controlled lab setting (Götz, 1964; Benzer, 1967; Heisenberg and Buchner, 1977). At the same time, the low cost, small size, wide availability, and ease of breeding have made flies ideal for use in educational and outreach settings, especially as the first or only hand-on introduction to genetics for many students (Harbottle et al., 2016). One important benefit of popularizing *Drosophila* methods for educational settings is that cutting-edge research can become directly relevant to the experience of the students. However, it is challenging to bring modern methods in animal behavior to teaching laboratories, since most setups developed for this purpose are built from custom components that are often quite expensive or difficult to obtain. Whereas just a few years ago, specialized components required custom machining or complex procurement, due to the surge of ‘desktop manufacturing’ and tools like 3D printers and laser cutters, together with increasing interest in citizen science and STEAM education, there is a push towards making complex laboratory setups widely accessible. Here we describe our efforts to optimize the cost and accessibility of a complete system for preparing and experimenting on flies using the preferred method in our laboratory — precise behavioral measurements for single, body-fixed (tethered) flies presented with controlled visual stimuli (Reiser and Dickinson, 2008; Dombeck and Reiser, 2012). From our prior experience, these experiments are a uniquely appealing entry point for teaching students about experimental neurobiology, for an introduction into laboratory instrumentation, and for hands-on exposure to quantitative animal behavior and the related opportunities for stimulus designs and data analyses. We hope that the low cost and accessibility of this system makes it suitable for a wide variety of laboratory settings, from summer courses to undergraduate and even high-school teaching labs.

In what follows we describe the motivation and goals of the project, then detail all of the components of the system, characterize the performance of the integrated setup, demonstrate its performance in measuring rather sophisticated aspects of visually guided behavior in flies, and finally estimate the cost of our systems. While we favor a modular, adaptable approach to instrumentation, we have endeavored to simplify the described system, so we mainly detail one specific setup, but throughout we describe some alternative solutions that we considered. The manuscript describes the system that we have built and used for data collection between November 2020-March 2021, and is thoroughly documented at https://reiserlab.github.io/Component-Designs/. We expect to continue making improvements to this setup and will post updates to that repository.

### 1.1. Motivation and Approach

The continual improvement of many commercial technologies comes as a direct result of massive, iterative efforts, by thousands of engineers, optimizing all aspects of the design of these products (consider that smart phones are not quite 15 years old). By comparison, even the most mature instruments used for collecting laboratory data are essentially bespoke prototypes benefiting from very few ‘generations’ of development. For that reason, many scientists prioritize designing their setups to combine high flexibility with precise control, which often requires using fairly expensive components capable of precision that far exceeds the requirements (often overestimated since never precisely specified) of any individual experiment. For the fly behavioral setup we have sought to optimize, we now benefit from several decades of methods development by many laboratories, which means we understand the requirements of this system rather well. Consequently, by eliminating unnecessary precision and flexibility, and taking advantage of desktop manufacturing tools, we could greatly simplify these setups and can now replicate them at much lower cost.

The setup we detail here was initially inspired by an invitation to run a hands-on training module at the *Drosophila* Neurobiology: Genes, Circuits & Behavior course at the Cold Spring Harbor Laboratory during the summer of 2019. We know that a front-of-class demonstration does not come close to the hands-on-experience of anesthetizing and tethering flies and then positioning them on a treadmill to observe walking behavior, but this requires many, independent setups. In preparation for the course, we began by attempting to replicate the typical (overly flexible and precise) walking fly-on-ball setups we favor in our lab, that are largely based on the systems we have previously described (Seelig et al., 2010; Strother et al., 2017). We focused on replacing the most expensive components, one-for-one, with less expensive commercial parts and some 3D printed components. At the time of the course we had converted a setup that would cost ~ $16,000 to replicate, to one that could be built for <$500. We assembled seven of these setups and succeeded in providing rigs to small groups of student who all learned to glue flies to pins and to position them on the treadmill. While this version was quite successful, we were unsure if it would simply be suitable for demonstrations, or whether it could fully replace our typical setups. In the past year we have continued to simplify and optimize the setup, with the goal of attempting to reproduce our gold-standard data set, the so-called optomotor response of walking flies (Götz and Wenking, 1973; Buchner, 1976), with its well-studied dependence on the spatial and temporal properties of the visual pattern. Due to the COVID-19 pandemic we did not return to the course during the summers of 2020 or 2021, but have continued to refine the setup, so that we now have a complete, full-featured, low-cost implementation of both a fly preparation setup and the experimental setup, which is described below. Component designs, including those we plan to implement in the near future will be shared at https://reiserlab.github.io/Component-Designs/.

## 2 MATERIAL AND METHODS

### 2.1. System Overview

We detail the major components of our system for preparing (tethering) and measuring the walking behavior of flies. The components of the experimental setup are shown in Figure 1 **a,b**. A single fly is tethered to a rod, which is mounted on a manipulator allowing for precise positioning of the animal along the three translational axes (all components color-coded; manipulator in blue, Figure 1 **a**). The fly is positioned on top of an air-supported sphere, which serves as an omnidirectional treadmill (sphere holder in green). The temperature near the fly can be regulated via a heater below the ball holder (in purple); a thermistor attached to the holder provides the measurements for closed-loop control. Visual stimuli are displayed on a tablet computer (in gray) and rotations of the ball in response to fly walking are captured by the camera (in red). The ball is illuminated by three LED fixtures (in yellow). Figure 1 **c** shows signal flow for the system, including a computer that runs the software for ball tracking (FicTrac (Moore et al., 2014)) as well as FlyFlix, the software we developed to generate stimuli and log responses.

**Figure 1.**
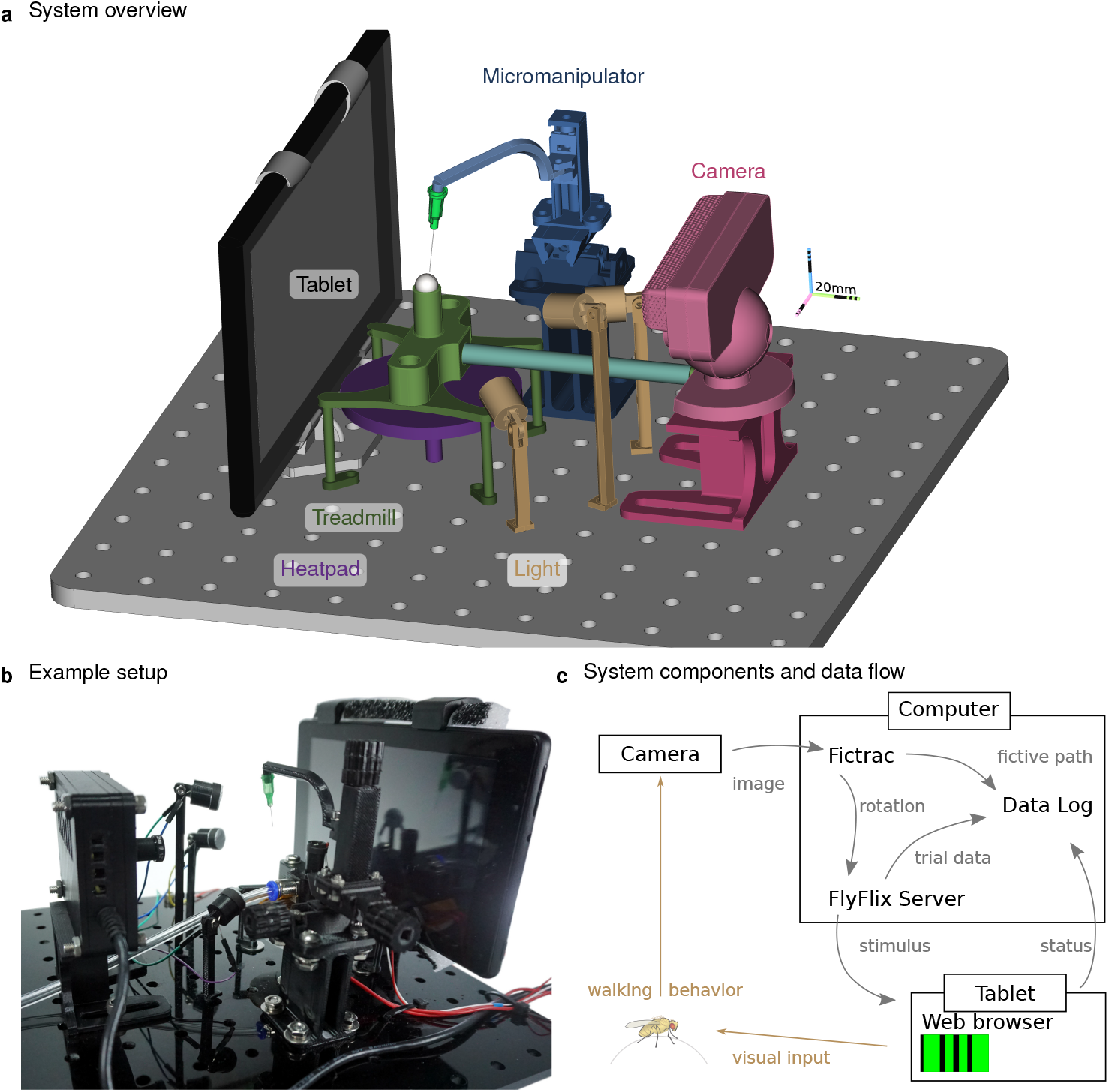
Inexpensive treadmill setup for walking fly experiments. The rendering in **a** highlights the major components. The fly is tethered to a thin syringe tip (light-green cone) and positioned and held in place with the micromanipulator (in blue) facing the tablet (gray) while walking on the treadmill sphere (in white). The treadmill holder (in green) floats the sphere on a steady stream of air supplied by the (light blue) tubing that pointing towards the camera (in red) used to track the sphere rotations. Adjustable lights (in yellow) illuminate the sphere. The temperature near the fly is controlled from below by the heatpad (in purple). All components are mounted on a breadboard laser-cut from an acrylic plate. **(b)** A photograph of the setup in the lab. **(c)** The flow of information between the major functional modules. For closed-loop experiments, ball rotations from FicTrac are routed to FlyFlix for on-line stimulus updates.

Gluing flies to a thin rod, a process referred to as *tethering,* can be straightforward, but requires a specialized setup that is not widely available or particularly well described in the literature. Therefore, we set out to simplify and document our solution here (Figure 2). A good tethering strategy must enable the precision required for positioning at the small scale of the fly body, as well as the mechanical robustness required to be manipulated by human hands. Essentially, a small fly needs to be carefully glued to an object that people can routinely move from one device to another. It is nearly impossible to tether a moving fly, and so flies must first be immobilized. While there are multiple ways to anesthetize flies, and CO2 is commonly used, this gas is known to have many effects on behavior that can last for hours. Instead we favor chilling flies, which causes insects to enter a chill coma due to a transient failure of neuromuscular function, from which they rapidly recover (Findsen et al., 2014). When chilled to temperatures close to freezing (Gibert and Huey, 2001), flies rapidly immobilize.

**Figure 2.**
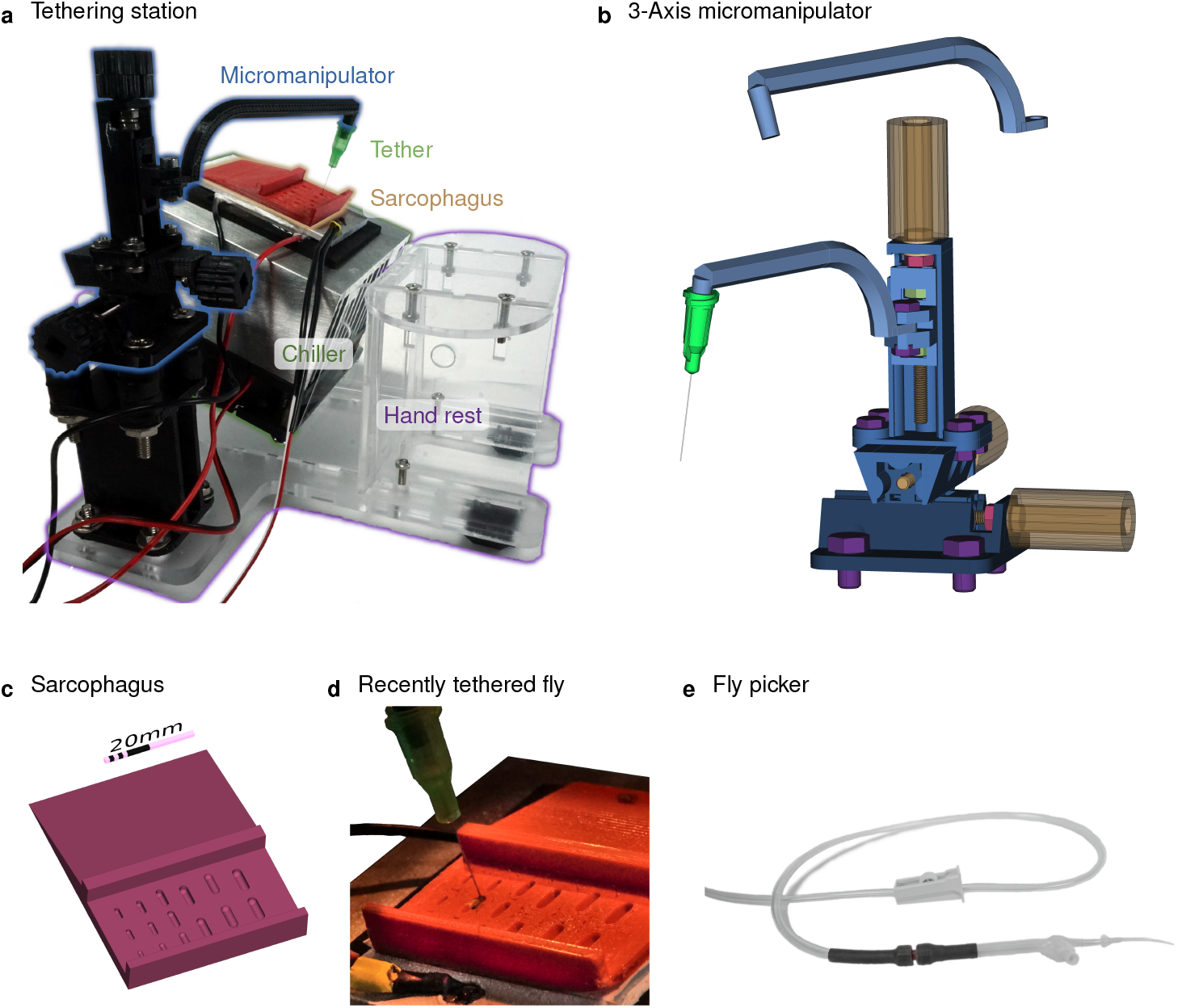
Chilled tethering station for preparing flies. **(a)** The major components highlighted in this photograph of the tethering station: micromanipulator (in black), sarcophagus (in red), Peltier-based chiller with heat-sink (silver and black), and a transparent laser-cut fixture with a hand rest. This setup is typically used under a dissecting microscope, and a thermistor is used to measure the top surface of the chiller for closed-loop temperature control. **(b)** A compact, 3-axis micromanipulator fabricated from 3D-printed parts and simple hardware components. Two different arms are used, one for the tethering station and the other for the experimental rig. For each axis, the rotating handle and screw are color-coded in yellow, the location of the screw is fixed in the outer rail via the red locking nut, and the green nuts within each carriage transfer the liner motion. The device is held together and mounted using the purple screws **(c)** A rendering of the sorting and mounting plate, containing a series of indentation, each referred to as a *sarcophagus,* of different dimensions for different animal sizes. Cold-immobilized flies are sorted on the top section of the plate, and single flies are positioned in one of the cavities for gluing to the tether. **(d)** A photograph of the plate mounted on top of the chiller with a temperature sensor (yellow tape) and a fly glued to the tether (a dispensing needle). **(e)** The fly picker used to move anesthetized flies. The picker uses suction controlled by the operator’s finger to pick up and deposit single flies.

We place vials of flies directly into an ice bucket for 5min (not shown). We then transfer the immobile flies to a cold, temperature-controlled platform, the ‘Chiller’ in Figure 2a. We tap out a small number (<5) of flies onto the sorting platform on the upper part of the red structure in Figure 2a. We use the fly picker (Figure 2e) to gently pick up (using suction) a selected fly (usually a large female) and deposit the fly into one of the semi-cylindrical indentations in the lower part of the platform (see Figure 2 **a, c, d**). On occasion it is possible to perfectly position a fly into this ‘sarcophagus,’ but typically the flies need to be aligned using a paintbrush before a small drop of glue is placed (with a fine wire, see Figure S5) towards the anterior side of fly’s notum, the dorsal surface of the thorax. We have designed a hand rest to support the user’s arm during these fine-scale manual steps (see Figure S4). We then use the three-axis linear stage Figure 2a,b to position the tether so as to just make contact with the glue. We use glue that rapidly cures upon illumination with short wavelength (ultraviolet, UV) light. Once cured, the fly can be lifted out of the sarcophagus and is ready to be used in experiments within a few minutes (Figure 2d).

We describe the construction of the tethering station in Section 2.2 and the experimental setup in Section 2.3. In Section 2.4 we detail how we used these components to run experiments.

### 2.2. Tethering station

For maximal user convenience, we recommend that the tethering station be physically separated from the experimental rig (as in Figure 2) and positioned under a dissecting microscope. However, even further cost and space reduction are possible if the tethering station and experimental setup are integrated into one unit (see Figure S3), making use of a single micromanipulator for both.

#### 2.2.1. Magnification

In our experience, every student can learn to prepare excellent flies for behavioral experiments with only a few sessions of practice. However, better results require learning to position flies so they are glued with approximate symmetry—in the center of the anterior notum and with minimal body rotation about the roll and yaw axes, and with the tether glued at 90° to the body long axis. This precision requires magnification. In our current setup, we position the tethering stage below a salvaged stereo microscope (Zeiss STEMI SV8). We have confirmed that flies can be tethered with alternative magnification methods such as a low-cost ‘toy’ USB microscope and a magnifying glass typically used for soldering electronics, however neither is as practical as a stereo microscope. In particular, we find that the instantaneous feedback of an all-optical system is ideal for mastering the hand-eye coordination required, while the delays in the low-cost digital microscope were quite challenging to work with. We recommend a stereo microscope with magnification of at least 10X (although 30X is even better) that can be lifted to support the height of the tethering station, which requires clearance of at least 170 mm.

#### 2.2.2. Tether

The tether provides the critical interface between the humans-scale and the fly-scale. We base the tethers in our laboratory on the original three-part design of Michael Dickinson (Lehmann and Dickinson, 1997). These parts are a connector, a 0.1mm diameter tungsten rod (that gets glued to the fly), and hypodermic tubing to connect the two parts. The rod and tubing are typically sold in longer units and need to be cut to length before the three parts can be assembled, and we require a special setup to assemble these components reliably. Also, the tethers can be easily bent and require regular repairs and replacement. Because of the laborious assembly, and other limitations of this design, we tested many alternative options that would be well suited to the needs of a teaching course.

In our walking setup, we obtained excellent results using unmodified blunt-tip dispensing needles. Dispensing needles with Luer adapters are widely available, manufactured to tight tolerances, inexpensive, and easy to handle. We selected 34 ga needles, featuring a stainless steal tube an outer diameter of 0.25 mm and 0.5 in (about 12.5 mm) in length. This is the finest needle size that is readily available from many vendors (e.g. AG-ABSS-99D0, Bstean, China). Due to this fine size, these dispensing needles are also suited for tethered flight experiments, but are easily bent and should be handled with some care. We use the inner sloped cone of the Luer lock to friction mount the tethers to our setup. We designed mount points on the arms of the Micromanipulator in the preparatory and experimental setup (Figure 2b). It is convenient to tether a number of flies one after the other and hold them until experiments are carried out (see Figure S5 i). We hold flies on a strip of upward facing screws (M4 or 8-32 will work) glued to a surface. We note that the Luer adapter is keyed with a pair of plastic tabs that can be used for alignment. We only use these as a visual aid, but this feature could facilitate more automated alignment in future setups.

#### 2.2.3. Glue and curing

To fix the tether to the fly thorax, we use resins that polymerize upon intense illumination, conveniently converting from liquid to solid within seconds. The standard glue used in our lab is KOA 300 (Kemxert, Poly-Lite, York, PA, USA), that requires UV (320 nm– 380 nm) light to cure. We typically use a commercial spot-curing lamp, such as the SpotCure-B (Kinetic Instruments Inc., Bethel, CT, USA), as they supply high intensity illumination (that cures the resin within seconds) and feature a convenient, audible timer that allows us to achieve consistent curing. Throughout the development of our setup, we tested different glue products and have confirmed that Bondic UV liquid plastic (Bondic USA, Niagara Falls, NY, USA) and Solarez Fly Tie UV Cure (Solarez Wahoo International, Vista, CA, USA), which are both widely available, work well as the tethering glue. We have a slight preference for the viscosity of the KOA 300 glue. Bondic and the Solarez thick formula appeared to be more viscous, while the Solarez flex formula (green package) is quite similar to KOA 300. The cost differences between these options are not significant, and we used KOA 300 in our experiments.

For teaching lab applications, we have used inexpensive UV-LED mini flashlights (basically a CR2032 battery with a single UV-LED) to cure KOA 300. Bondic and Solarez are available in packages with battery-operated curing lights that work well for our application. In the near future, we plan to integrate automatically timed UV curing lights into the tethering setup.

#### 2.2.4. Cooling

When cooled below 4 °C *Drosophila* are rapidly and reversibly immobilized (Gibert and Huey, 2001), which makes it convenient to align and tether the flies, and also to perform further surgeries and treatments if desired. While the flies can be chilled on a metal stage mounted on ice (or a frozen gel ice pack), a temperature-controlled thermoelectric cooler provides a more compact and stable solution. Using the Peltier effect, a powered thermoelectric cooler moves heat from one side of the device to the other, but to cool one side stably below room temperature, heat from the hot side must be effectively carried away. In our lab, we use a recirculating water chiller to pump water through a liquid-cooled Peltier assembly. This is quite reliable, but is expensive and cumbersome (substantial tubing is required), and occasionally very messy.

For our optimized setup we use an integrated, low-cost, air-cooled Peltier assembly that consists of a 40 mm × 40 mm thermoelectric module mounted between a 40 mm × 60 mm aluminum plate and a 90 mm × 90 mm aluminum heat sink with a fan (Adafruit Industries). We find that when powered with a 12 V 6 A supply, the top aluminum plate reaches temperatures below 0 °C while operating under typical ambient room temperatures. This confirms that the device can adequately cool flies. In order to provide a consistent temperature above freezing, we implemented closed-loop temperature control using a W1209 module (multiple vendors, e.g. MOD-78, ProtoSupplies, Lake Stevens, WA, USA) that can regulate an electric load up to 10 A based on input from a 10 kΩ NTC thermistor attached to the top side of the chiller’s aluminum plate (see Figure 2 **d**). We mount the chiller at 20° towards the experimenter, shown in Figure 2 **a**. This angle provides good airflow for cooling the module, while pushing air away from the experimenter so as not to blow flies off of the tethering station. By pitching the platform we ensure that flies will always be visible from above while being inspected, aligned, and tethered (both the fly and tip of the tether are seen throughout). For the simplest setup, we angled the chiller by extending two screws at the corners of the fan attached to the heat sink (see Figure S3). Instead, the integrated setup shown in Figure 2 **a** is assembled from laser-cut acrylic sheets. This design also includes mounting holes for the micromanipulator and a hand rest (CAD files in the accompanying repository).

#### 2.2.5. Sarcophagus

To position, hold, and sometimes dissect or manipulate cold-anaesthetised flies during tethering, we typically use a movable cylindrical cavity machined from solid brass. This design is affectionately referred to as a *sarcophagus* and based on the original design of Karl Götz (Max Planck Institute for Biological Cybernetics) from the 1960s. The most important feature of the cavity is that it should be smooth and slightly larger than a fly, since fly legs can be easily caught on sharp edges. Beyond this detail, many aspects of the elaborate Götz design are not required for routine tethering of flies for walking experiments. For the optimized tethering stage, we tested various 3D-printed sarcophagus components. We find that parts printed from different materials, including resin, ABS, and TPA, all worked similarly well.

The example plate in Figure 2 **a,d** is made from red ABS. 3D printing allowed us to place cavities of different sizes on a single plate, to accommodate experimenter preference for size and depth, as well as to readily support tethering insects of differing sizes. We designed an inclined sorting area on the top section of of the plate, such that different regions are made of different depth of material, creating different temperature zones. We sanded the bottom side of the sarcophagus plate and mounted it to the aluminum plate of the chiller with thermal adhesive tape. We find that setting the Chiller to a nominal temperature of 2 °C works well for our setup. Flies remained immobilized while on the plate, but started moving within seconds off the plate. The temperature setting may need to be adjusted for different setups.

#### 2.2.6. Micromanipulator

For precisely and stably positioning the tether near the fly for gluing, we typically use industrial-grade three-axis linear stages with a probe-clamp (from Thorlabs or Siskiyou). Because these devices are used for microscopic manipulation, they are referred to as *micromanipulators.* We use a second micromanipulator to position and then hold the fly on the treadmill of the experimental setup. These manipulators are essential components of reliable setups, but the commercial components we use are too expensive ($500 or more) for the purposes of a teaching course. In exploring alternative manipulators, we tested lower-cost, three-axis manipulators ($100, e.g. LD40-LM, multiple manufactures available through Aliexpress, China) and find them to be a suitable replacement for linear stages from lab suppliers (see Figure S4). However, we were interested in exploring even lower cost options, and evaluated several 3D-printed alternatives, including the micromanipulator design from Open Labware (Baden et al., 2015; Chagas et al., 2017). We find this design to be quite workable, but the footprint was somewhat challenging to incorporate into our setup. Based on these explorations, we proceeded to design our own micromanipulator, optimized for simplicity and cost, and with a compact footprint.

Our three-axis micromanipulator design (Figure 2 **b**) is assembled from nine 3D-printed parts and standard screws. For each axis, an outer rail surrounds the carriage on three sides. Each rail features a screw held in place by a locking nut (red in Figure 2 **b**). Turning each yellow knob with the attached red screw moves the corresponding green nut, and with it the carriage. The arrangement of the three axes allows translational movement in any direction by up to 20 mm. We printed the parts from ABS on a F-170 (Stratasys Ltd, Eden Prairie, MN, USA). This design requires slightly tighter tolerances than can be relied on from the printer, so we sanded and smoothed the outer faces of the carriages with 200 grit sanding paper until they would slide into the rail. The rails did not require post-processing. Even though this sanding can take up to 30 min, we find this advantageous as it allows us to produce a close fit across material and printers, and thus high accuracy movement, without adapting the design. We also recommend applying a plastic lubricant to the rails (e.g. Dry-Film Lube, WD-40, San Diego, CA, USA) to increase smoothness of movement. 3D printed (or laser cut) knobs are glued to the screw heads to allow comfortable handling of the micro manipulator. The bottom rail has additional mounting holes to attach the manipulator securely to the baseplate.

We have designed two arms for mounting to the z-axis carriage, shown in Figure 2 **b**. The floating arm is used for tethering flies, is slightly longer, and holds the tether (by a friction fit) at 20° inclined from vertical, to match the angle at which the heat sink is mounted. The arm shown as fixed to carriage holds the tether at a 10° angle in the opposite direction, and is used for the experimental setup. The orientation and function of the arms can be rapidly adapted to specific applications. The micromanipulator assembled from 3D printed parts is a low-cost functional substitute, but it does not replicate the properties of a commercial linear stage. In particular, the plastic parts are somewhat compliant and cannot be used with heavy loads.

#### 2.2.7. Fly picker

Single flies need to be moved and carefully positioned on the tethering platform. It is possible to do this with forceps, but we do not recommend picking up flies destined for behavioral experiments by either their legs or wings. In our laboratory we use a commercial vacuum pump and wand with a very fine tip that is typically used for handling tiny electronic components during assembly. With such a device it is possible to very gently lift a fly and quickly deposit them into the sarcophagus in nearly the ideal position for tethering. One alternative would be to fit a standard lab aspirator (or pooter) to use a fine tip. However, we find the hand-held vacuum approach to be rather convenient and so we have fashioned a version from standard components (Figure 2 **e**). We use a plastic transfer pipette with the bulb end cut off and replaced by a tubing connector (we used Luer locks connectors, but any tight connection would work). This connection is further strengthened with heat shrink tubing. We connect the tubing to our available lab vacuum, but this can be replaced by suction pumps or other sources of negative pressure (Baden et al., 2015). We control the the suction from the picker by a roller clamp on the tubing. We cut a hole in the side of the pipette and glued in another adapter with a flat surface. When covering this stub with a finger, the suction at the tip substantially increases. When the finger is removed, the fly is released. Since the opening at the tip of the transfer pipette is too wide for a *Drosophila,* we added a one-way tip (F1732011 Pipetman Expert Tips EL10ST, Gilson, Middleton, WI, USA) as in Figure 2 **h** or piece of thin heat shrinking tube in Figure S5g. By bending a paper-clip to a desired angle and using a heat gun on the shrink tube and plastic pipette tip, we bent the tip in an angle that allowed more convenient use. We find that the pipette tip with an inner diameter of 0.25 mm and an outer diameter of 0.65 mm allows for convenient manipulation of flies. Flies are incredibly robust, but we recommend adjusting the pressure (via the clamp) to just above the threshold for reliably lifting flies.

### 2.3. Experimental Setup

Here we detail the major components already introduced for the walking fly on a ball setup shown in Figure 1.

#### 2.3.1. Baseplate

Many lab setups are built on solid aluminium breadboards with threaded mounting holes from specialized lab equipment manufacturers. They are very stable and can be flexibly used for many purposes. In place of these boards we use a 300 mm × 300 mm × 10 mm acrylic board into which we cut 144 holes of 6.35 mm diameter in a 12 x 12 grid with 1 in (2.54 mm) spacing using a laser cutter. To further simplify the design, we opted not to tap threads into the holes. Several components, such as our LED lamps, can be flexibly positioned using a friction fit. Parts that require stable fixation can be mounted with nuts and screws. We use adhesive rubber feet at the corners to lift the baseplate and add some vibration damping. This baseplate could be further simplified to the minimal size and number of mounting holes required to fit the components in the setup, but the additional holes allow for future extensions to the apparatus. This simple design is both light and stable, ideal for carrying to teaching labs and outreach events.

#### 2.3.2. Micromanipulator

To walk with a typical gait the fly needs to be positioned ~0.4 mm from the surface of the sphere, and aligned to the center of the ball (see Figure S5 j). We use the identical micromanipulator, of our own design, described in section 2.2.6, with the arm that positions a fly so they are walking at 10°, or slightly ‘uphill’ – based on observations made in our lab, which appears to improve walking performance (personal communication, Shiuan-Tze Wu). The tether is friction mounted onto the arm and can be gently rotated so as to manually align the long axis of the fly towards the screen.

#### 2.3.3. Treadmill

The omnidirectional treadmill consists of a stem that holds an air-supported sphere. Our simplified, 3D printed design for the sphere holder is a direct adaptation of the design from Seelig et al. (2010), which was custom-machined out of aluminum. The original design made use of a straight inner shaft for airflow to simplify the machining process, but this limitation does not apply to 3D printing. We implemented a more compact design where air enters via tubing with a 90° angle to the sphere-supporting air column, as shown in Figure 1 **a,b**. In addition, we provide CAD files for alternative designs, including for different ball sizes, in the accompanying repository.

While some 3D printing methods will produce a solid, airtight part, most printers that build up parts by fusing filament in layers may result in parts that are not airtight and will allow air to escape. Rather than require specific printing methods, we find that quite satisfactory performance can be achieved with simple post-processing. Applying acetone to the surface of the printed part seals the holes. We find that sealing only the outer surface works well, while applying solvents to the thin inner tubing could result in clogging the air stream (which can be drilled out).

The flow-rate of the air needs to be controlled: if too low, the ball won’t reliably float, and if too high, the ball will be much less stable (or fly off). In the fly-on-ball setup of Seelig et al. (2010), the airflow is fed by pressurized air which is regulated by a commercial mass flow controller. We find that with no loss of performance an inexpensive flowmeter can be used instead, but it is important to select one that allows fine control over the appropriate range of airflow. For example, VFA-22 (Dwyer, Michigan City, IN, USA) with a maximum of 1 Lmin^-1^ works well. In practice, we adjust the flow rate by visually inspecting a walking fly on the ball. In place of a pressurized air supply, we have tested an aquarium-style air pump with a maximum flow rate of 1.8 L min^-1^. We find that inexpensive pumps induce some vibrations in the ball and are continuing to investigate the ideal substitute for wall-supplied pressurized air.

#### 2.3.4. Spheres

The sphere of the treadmill is the only moving part during the walking experiment and is therefore crucial for good measurements of behavior. The sphere needs to be nearly perfect in shape but not too smooth, light enough to float and be spun by the fly, but not so light that flies can pick it up, and with low rotational inertia to enable mostly unrestricted walking by flies. We have tested many alternatives, but our preferred standard sphere is still based on the method of Seelig et al. (2010), where the spheres are cut from foam with either a file (design available at https://wiki.janelia.org/wiki/display/flyfizz) or by a CNC machine. We find that flies walk best on a sphere cut from Last-A-Foam FR-7120 (General Plastics Manufacturing Company, Tacoma, WA, USA) to a diameter of 9 mm (density of 320 kgm^-3^, sphere weighs approx. 0.12g). For optical tracking of the sphere rotation with FicTrac (Moore et al., 2014), we paint this NIR-reflective foam with BLK3.0 paint (Stuart Semple studio, Dorset, UK), which we find to be less NIR-reflective than a black permanent marker, and thus produces high contrast features. We continue to test alternative sphere materials that will be more readily available than a hand-filed foam ball. The results will be documented in the accompanying project repository.

#### 2.3.5. Sphere tracking camera

In tethered walking experiments the flies are fixed in space, however their intended locomotion, as if walking on an infinite virtual plane, can be estimated through the rotation of the sphere they are turning. Several methods have been developed for measuring relative rotations of the ball, for example through optical mice sensors or via optical flow calculated with camera-based systems (Lott et al., 2007; Seelig et al., 2010; Vishniakou et al., 2019). Under ideal circumstances, these relative measurements can be calibrated for excellent accuracy, but these systems can be quite sensitive to the uniformity of the lighting and focus of the sensors, etc. By estimating the absolute position of the sphere in every frame, the tracking software FicTrac (Moore et al., 2014) is an exciting alternative approach that offers several advantages.

To use FicTrac, the sphere must be painted with a high-contrast pattern that looks unique at any position. Fictrac then maps individual camera frames to a previously constructed template of the sphere’s pattern to estimate the instantaneous rotation of the sphere. The animal’s virtual trajecotry can be recosntructed from the frame-by-frame estimates of the sphere’s position. The software works best with sharp edges and high contrast, so Moore et al. (2014) suggest to avoid motion blur, by imaging with high frame rates. FicTrac supports industrial cameras from Flir and Basler, as well as images through OpenCV (https://opencv.org/), a library for real-time machine vision with extensive support for a variety of cameras. As FicTrac operates on a high-contrast single color channel image, downsampled to a resolution of 60 px × 60 px, we realize that the ideal low-cost camera for this application should support a low-resolution, high frame rate mode (a combination of requirements that are nearly the opposite of most inexpensive camera sensors). We found that the PlayStation Eye camera (Sony Entertainment Corp.), developed as an input controller for action games, is an excellent solution. Using open-source drivers for the low-latency integrated video processor, we obtain access to a stable video stream of 187 fps at a resolution of 320 px × 240 px. The camera sensor OV7720 (OmniVision Technologies) was developed for low-light operations, and we found that the sensor is quite sensitive to infrared illumination (once the filter, attached to the lens housing, is removed). To effectively use this camera, we have modified the body for easier mounting and to accept S-mount lenses, as shown in Figures S1 and S2. For reliable imaging of the sphere at a working distance of 10.5 cm we mount a macro lens with 25 mm focal length and an intermediate (fixed) aperture (see Figure 1 **b** and Figures S1 and S2).

The PS Eye camera is our preferred low cost solution, but due to the modularity of our setup and FicTrac’s support for many cameras through OpenCV, many other cameras with similar properties should work. To ensure that our system works reliably on readily available PCs, all tests were run and data were collected while running FicTrac on an older model, multi-core x86-64 system with a maximum frequency of 3 GHz and a hard disk drive running Lubuntu 20.4 LTS. This PC was powerful enough to run two FicTrac instances as well as FlyFlix, the software we developed for stimulus presentation, experiment control, and data logging, and so we expect that most PCs will be able to run these experiments.

#### 2.3.6. Lighting

An important consideration for measuring visually guided behaviors is to use illumination that minimally interferes with the animal’s vision. The most practical solution is to use near-infrared (NIR) illumination since fly vision is insensitive to these longer wavelengths (Sharkey et al., 2020), but most camera sensors measure it well. As we operate our camera at high frame rates and with an intermediate aperture lens, intense illumination is essential for reliable sphere tracking, yet the light cannot be so intense as to saturate regions of the image (due to the limited dynamic range of any camera).

To achieve strong, but diffuse NIR illumination, we use three generic 840 nm LEDs placed between the camera and the treadmill, pointing towards the sphere. We designed compact 3D-printed housings that allow flexible positioning of the light sources at the top of posts that are friction fit into the holes on the baseboard, as shown in Figure 1 **a,b**. We used pieces of a plastic bag as a diffuser in front of each lamp, attached with heat shrink tubing. For our setup, we used a 5 V power supply together with a 470 Ω current-limiting resistor. With these lamps in place we adjust the lights until we obtain images of the sphere that are bright, yet evenly lit. In our standard setup (our display set to 25% brightness), we do not need to place a visible light blocking filter on the camera, although this could be required for robust ball tracking with other displays.

#### 2.3.7. Heatpad for temperature control

We typically run experiments in rooms that are climate-controlled for the comfort of humans, yet these conditions are often not ideal for flies. It is often convenient to increase the temperature of the walking fly, both since flies walk more often and faster at elevated temperatures (Soto-Padilla et al., 2018) and also for using temperature-dependent genetic reagents such as Shi^ts1^ or TrpA1 (Owald et al., 2015). To inexpensively support warming the fly, we installed a resistive heater pad underneath the sphere holder, controlled by a second XH-W1209 temperature controller (also see Section 2.2.4). We attached a thermistor to the sphere holder as close to the animal as possible. The actual temperature at the animal position might be slightly different (and should be verified if critical), and we consequently refer to the the XH-W1209 setting as the *target temperature.* For the experiments detailed below we use a target temperature of 32 °C.

#### 2.3.8. Display

A surprisingly wide range of visual stimulus delivery strategies have been used for insect behavioral neuroscience: from motor operated moving objects like patterned drums, to projectors and computer monitors, to custom-made LED displays (Palermo and Theobald, 2019; Kócsi et al., 2020; Kaushik et al., 2020). In our lab, we typically use custom-made, modular LED displays configured as cylinders around the animals, to deliver stimuli with excellent temporal precision (Reiser and Dickinson (2008) and future developments documented at https:/reiserlab.github.io/Modular-LED-Display/). We have not yet succeeded at producing an inexpensive, widely available display using LEDs, and so we explored other options.

For the inexpensive treadmill setup, we used a widely available tablet computer with an in-plane switching (IPS) liquid-crystal display (LCD), an Amazon Fire 7 with a nominal screen size of 7 in. We display visual patterns through a web browser. To allow replication across devices we used Mozilla Firefox instead of the pre-installed browser. We installed the most recent versions of Firefox and kept the Android 9 based Fire OS updated with the latest release (most recently Firefox 86.1.x and Fire OS-7.3.x). We manually set the display brightness to 25 %. The tablet was connected to a USB power supply and connected to a local WiFi network during all experiments. To our knowledge, inexpensive tablets have not been used to test detailed behavioral responses of flies to moving stimuli, and so we evaluated both the technical performance of the display system (Figure 4) as well as the behavioral responses of flies to tablet-displayed motion stimuli (Figure 5). Tablets featuring IPS displays with 60 Hz refresh rate are the most widely available inexpensive option. It will be interesting to reevaluate new display technologies (such as OLED) with higher refresh rates as these become less expensive. Our existing system could be rapidly adapted to using a student’s personal smartphone instead of a tablet, further reducing cost (and probably distractions) in teaching environments.

#### 2.3.9. FlyFlix

By designing our inexpensive treadmill setup around a network-connected tablet as the visual display, we remove the need for any specialized IO devices for data acquisition or graphics cards for stimulus generation, but we needed to develop software we call FlyFlix, to control experiments, generate stimuli, and log data. Figure 1 **c** shows a simplified flow of information through the experimental setup. Our display connects through the web browser to the local URL of the FlyFlix server. Upon connection, the web server delivers the most recent version of the FlyFlix client software (written in JavaScript) as an HTML5 web page. The implementation follows an event-based approach with minimal dependencies between client and server, so that any device capable of displaying an HTML5 website can act as a client without prior installation of client software. We have verified that smartphones and computer monitors can be used to display the stimuli, but all results reported in this paper are from experiments using the tablet described in Section 2.3.8.

The FlyFlix client and server communicate over a bidirectional, low-latency WebSocket connection. The server can deliver different experiments at specific URLs, or different views on the same experiment to different displays (a feature not used in our standard setup). Once the client connects to the server, it shows a “Fullscreen” and a “Start” button, the first changes the client to a full screen mode, while the seconds sends the request to the server to start the experiment. Once the protocol finished or the WebSocket connection is interrupted, the FlyFlix client displays a button to “Reconnect” to the server. Once the client starts the experiment, the server generates a set of trials based on the pre-specified configuration. The FlyFlix client renders a scene based on its local representation of the stimulus. The server sends updated parameters to change the representation and the client continuously reports back the actual state of the rendered stimulus. This bidirectional communication happens throughout the experiments with time-stamped messages. The FlyFlix server was implemented in python-3.9 using the Flask-1.1.2 web framework. Bidirectional communication from the server-side is based on Flask-SocketIO-5.x with concurrent networking through Eventlet-0.30. The client is implemented in JavaScript and uses two external libraries: Socket.IO-3.1 for the communication and Three.js-r124 for rendering the visual stimuli.

For our ‘gold-standard’ visually guided behavioral experiments, we wanted to present moving grating patterns composed of vertical bars across the display. Depending on the condition, we move different patterns at different speeds through the frontal visual field of the fly. Using the 3D graphics library Three.js, these stimuli are represented as segments of a cylinder surrounding A virtual camera. These segments are defined from a basic material that emits color but is not affected by virtual lighting. The virtual cylinder is 305 mm in diameter, matched to the size of a typical LED arenas used in our lab. The height of the cylinder exceeds the size of the virtual camera frame. The virtual camera rendering accounts for the physical distance between the animal and display (35 mm), and has the effect of correcting the displayed size of the cylinder segments so that they span an equivalent azimuthal size from the fly’s point of view, but span a different physical size on the display (illustrated in Figure 3**b** for a grating made up of 10° bars). In the current implementation, we move the bars across the display by rotating the virtual camera, but we are updating this to support more complex closed-loop conditions, where the virtual orientation of the fly and the position of stimuli will be controlled independently.

**Figure 3.**
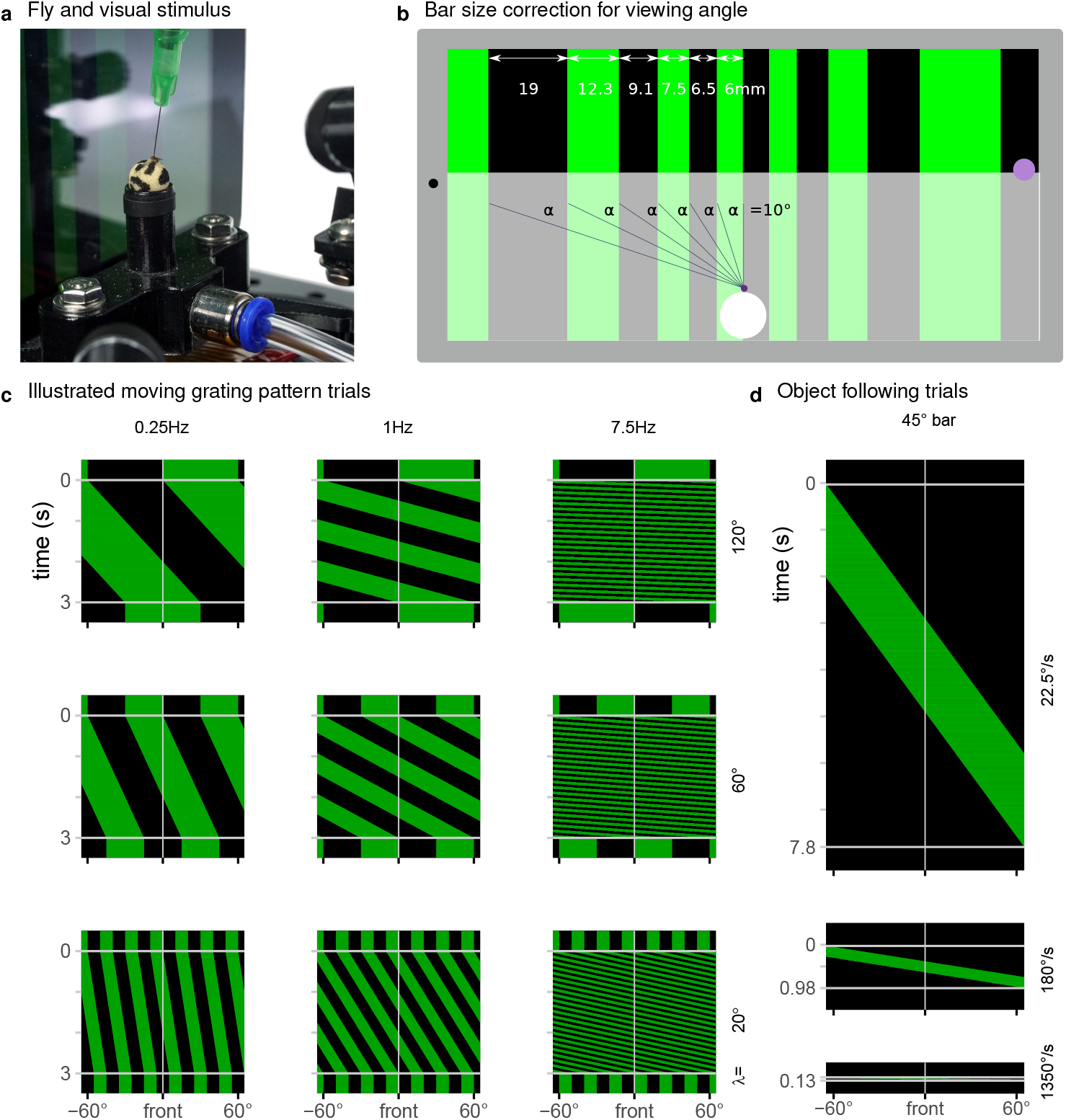
Display used to present a range of visual stimuli. **(a)** In our typical experiment, a tethered fly walked on an air supported foam sphere. The fly faced a mounted tablet computer that displayed a moving grating pattern. **(b)** The FlyFlix software renders a virtual scene that simulates a cylindrical display onto the flat screen. The azimuthal span of each bar within a grating pattern is scaled to correct for the viewing angle—even though the dark bar on the left is three times as wide as the bright one in the center, they both span 10° from the perspective of the fly positioned 35 mm in front of the display. The purple spot on the right marks the point of light measurement used for Figure 4 **a-c**. **(c)** Space-time representations of the display during trials showing a moving grating pattern. Each row of these images represents one horizontal slice through the displayed pattern, at any point in time. For these 3 s trials showing moving patterns with different spatial and temporal frequencies, clock-wise motion appears as space-time tilts that go down and to the left. **(d)** Representation of displayed screen content during object following conditions for clockwise movement with the indicated speeds.

Experiments are generated and controlled by the FlyFlix server. Depending on the experimental condition, the server asynchronously sends parameters describing the virtual scene to the client. These parameters primarily concern the scene layout and rotational speed, orientation, and maximum refresh rate of the camera. The client continuously renders new frames based on these parameters. This decouples the timing of server and client—the server does not need to consider the capabilities of the client such as the screen refresh rate, but instead communicates changes to the virtual world in real-time. Similarly the client is independent from the server—if there is a lag in the communication from the server to the client, the client does not have to wait for instructions but can instead render frames based on the previously communicated parameters. The client sends its time-stamped state to the server, and the FlyFlix server logs these together with its own time-stamped status to a file. We characterize the performance of this system and the network latency in Figure 4.

### 2.4. Experimental Protocol

To validate this new experimental setup we wanted to measure flies carrying out a well-studied visually guided behavior, the so-called syn-directional optomotor response, in which the flies steer, by turning, in the direction of a rotating visual pattern (Götz and Wenking, 1973; Seelig et al., 2010; Strother et al., 2017; Creamer et al., 2018). We recorded responses to open-loop stimulus presentations (in which the response of flies is measured, but not used to control the trajectory of the stimulus) of a periodic grating patterns moving at one of multiple speeds (mapping out the temporal frequency dependence) and a series of patterns composed of different grating periods (to map out the dependence on the spatial frequency). In addition, we also measured object-following behavior, by recording turning responses to single sweeps of bright or dark bars moving at different velocities.

For the temporal frequency tuning (14 conditions total), we showed grating with spatial period of *λ* = 90° composed of pairs of alternating 45° bright and dark bars. The periodic pattern moves either clockwise or counterclockwise with one of seven angular speeds *(ω* = 22.5, 90, 180, 360, 675, 1350 and 2700 °s^-1^). For a periodic pattern the temporal frequency is the angular speed divided by the spatial period *(ω/λ)* and so the tested conditions include 0.25, 1, 2, 4, 7.5, 15 and 30 Hz. For the spatial frequency tuning (14 conditions) we tested motion of gratings with one of 7 spatial periods (*λ* = 5°, 10°, 20°, 30°, 60°, 90° and 120°) all at a temporal frequency of 7.5 Hz, spanning angular velocities between 37.5 ° s^-1^ and 900 ° s^-1^. At the beginning of each trial, the initial position of the pattern is shown stationary for 500 ms, then moved for 3 s and shown stationary again for another 500 ms. Some examples of the patterns displayed in these conditions are shown in Figure 3 **c**.

In the object following conditions, either a bright or dark vertical 45° bar moves across the screen, exactly once at one of 6 angular speeds (*ω* = 22.5, 90, 180, 360, 675 and 1350 ° s^-1^) in either the clockwise or counterclockwise direction. Consequently these trials are have different length between 0.13 s and 7.8 s. During the 500 ms pre- and post-trial period, the screen is either fully dark for the bright object condition or fully bright for the dark bar. Diagrams illustrating these conditions are shown in Figure 3 **d**.

In our protocol, the open-loop conditions were interleaved with 3s closed-loop trials, where the fly’s turning controlled the position of the stimulus. We tested a variety of closed-loop conditions (data from these trials are somewhat ambiguous, and are not shown). For technical verification we extended closed-loop trials to a length of 30 s. These trials are also bracketed between 500 ms pre- and post-trials.

Within an experiment, each set of conditions was presented as randomly ordered blocks. The blocks were then repeated 6 times. We performed two separate experiments for the temporal and spatial frequency mapping data. At the beginning of each protocol there is a delay of 10 s to allow the experimenter to shield the experimental setup from the environment with a box, if desired. The behavioral results are presented in Figure 5.

**Figure 4.**
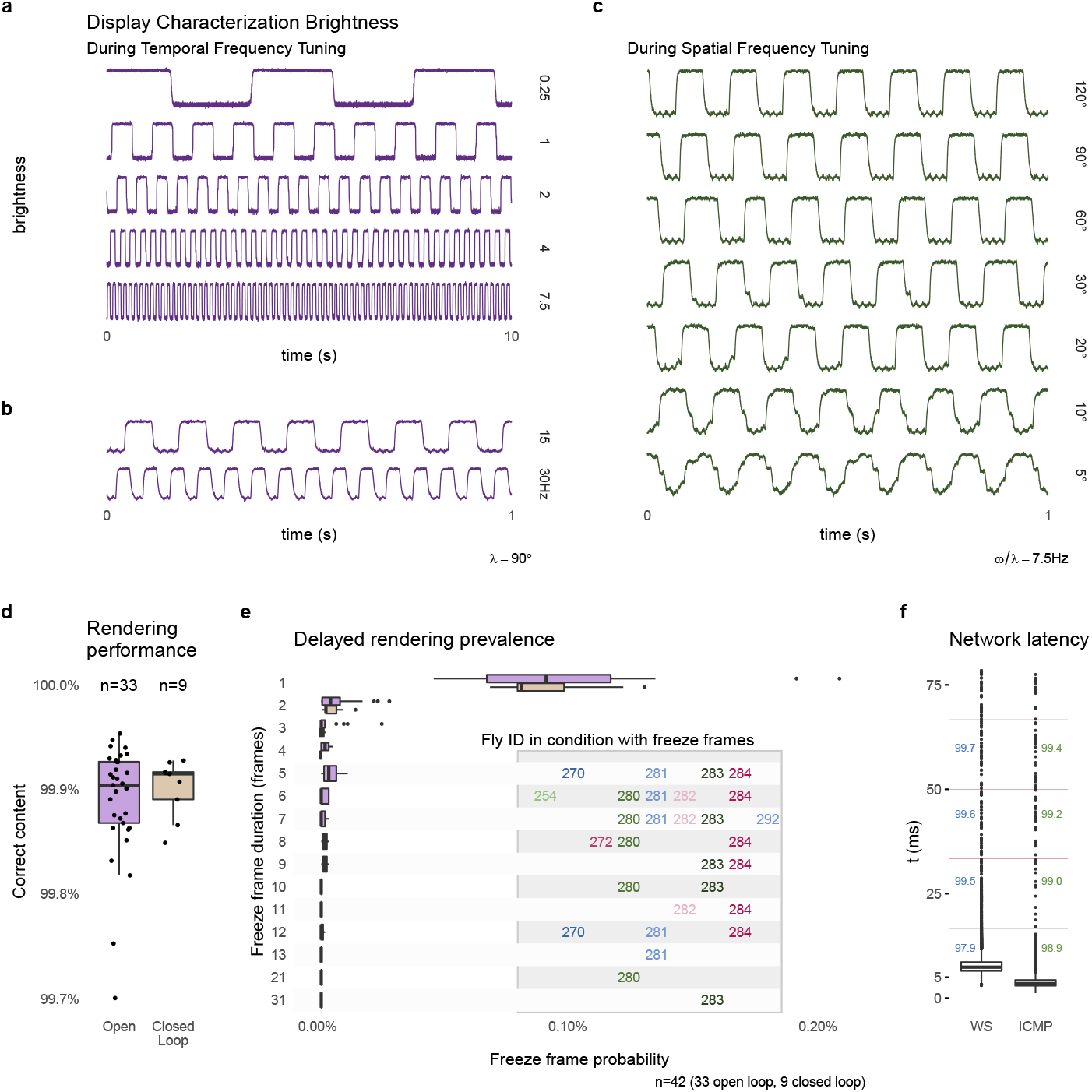
Technical performance of the experimental setup. Changes in the display brightness were measured at the approximate location marked in Figure 3 **b**, during presentation of moving grating patterns. The measurements show a regular pattern that changes at the expected temporal frequency (**a,b**) and at the same temporal frequency of 7.5 Hz during conditions showing different spatial frequency grating motion (**c**). The apparent filtering for the patterns with the finest bars is due averaging by the sensor. **(d)** The percentage of frames that are rendered correctly and within the expected time interval (see text for further details) during open-loop and closed-loop experiments. **(e)** Details of the frames that were not rendered within 1 frame interval. Most were delayed by a single frame, however, a very small number of longer delays occurred, but all from the same experiments. **(f)** The measured network latency for a round trip message between FlyFlix server and client through WebSocket compared to a network ping (ICMP). The numbers mark the percentage of round trips that would arrive within the 1^st^, 2^nd^, 3^rd^, or 4^th^ ~ 17 ms display frame (indicated by the vertical lines). Box plots show the first and third quartile for the box, median for the center line, the whisker extend to 1.5 of the inter-quartile range (IQR). d shows all data points, e and f only the outliers as individual points.

**Figure 5.**
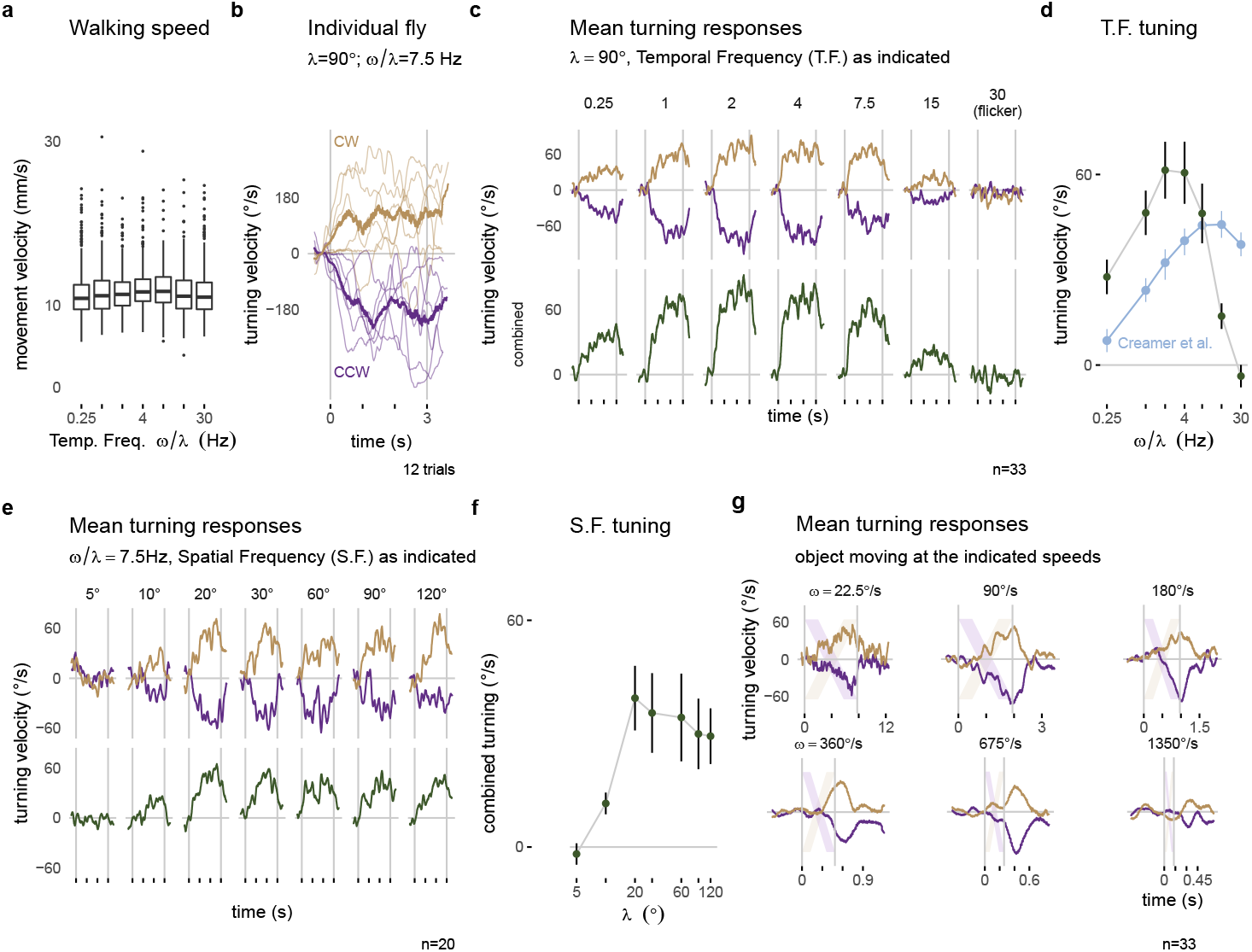
Visually guided turning behaviors measured with the optimized setup. Panels along the top row show walking behavior in response to a moving grating pattern with a spatial period of *λ* = 90°. **(a)** The forward walking speed of flies across the different conditions (n=33). **(b)** The turning response of a single fly to multiple presentations of 3s rotational stimuli moving clockwise (CCW, yellow) and counterclockwise (CCW, purple). Single trial responses are plotted as thin lines and the mean response across trials is in the thicker line. **(c)** The mean turning response across all 33 flies: the top row shows the responses to each direction of motion and the combined responses are plotted below. **(d)** The combined response, averaged during the period of stimulus presentation, is summarized as a tuning curve. For comparison the data in blue is extracted from similar experiments on a different setup, from Figure 1H of Creamer et al. (2018). The mean turning responses for a series of gratings with different spatial periods are plotted in **e** and summarized as a tuning curve in **f**. **(g)** The mean turning responses to presentations of a single 45° bar sweeping across the display. The lightly colored stripes represent the expected position of the bar on the display. The boxplot in a uses first and third quartile to span the box, 1.5 oIQR for the whiskers and outliers are plotted as individual points. The error bar in d, f are plotted as mean ± SEM.

### 2.5. Fly preparation

For behavioral experiments, we used the Dickinson Lab wild-type strain of *Drosophila melanogaster,* often referred to as DL flies. The flies were reared on standard cornmeal agar food at 21 °C and 50 % humidity. The speed tuning (Figure 5 **a–d**) and object tracking (Figure 5 **g**) experiment was conducted on 33 flies 4-7 days post-eclosion. Of these flies, 21 were male and 12 were female. 27 were raised in a 12:12 light-dark cycle and 6 were raised in a 16:8 light-dark cycle. The spatial frequency tuning experiments (Figure 5 **e**) were conducted on 7 flies (3 males and 4 females) raised in a 16:8 light-dark cycle.

Prior to tethering, we moved groups of approximately four flies from the fly vial to a 5 ml ‘Falcon’-style tube (12 mm wide) using a transfer funnel of our design (see Figure S5 (a, b)). After keeping the tube on ice for 5 min, the flies are immobilized and were gently tapped out onto the sorting plate of the tethering station. Individual flies were moved into the sarcophagus with the fly picker, aligned with a paint brush, had glue applied, and then the tether was positioned against and bonded to the fly with UV light curing as described in section 2.2. The chilling of the flies was kept brief, so that no flies were chilled for longer than 23 min. After lifting out of the sarcophagus, tethered flies were placed upside down in a holding area for at least 30 min between tethering and the start of the experiment. Flies were provided with a piece of 5 mm × 5 mm tissue, which they readily manipulate with their legs.

### 2.6. Data analysis

FlyFlix and FicTrac store the data in rectangular data files. While FicTrac follows a tidy data format, FlyFlix uses a key-value based long format. We use a custom python script to load these different data formats into a consistent SQLite database. Data analysis and plotting is done in R-4.x using the tidyverse-1.3 packages including ggplot-3.3.

Flies were presented with paired visual stimuli that moved in both the clockwise and counterclockwise direction. We recorded the ball rotation via FicTrac. Out of the 25 recorded variables, we used the ‘animal’s heading direction (lab)’ to estimate intended body yaw rotations and the ‘animal movement speed’ for the walking velocity. We used ‘delta timestamp’ to convert frame-based differences into time-based rotational velocity and the physical diameter of the sphere to calculate the movement velocity (Figure 5 **a, b**). For time series data, the average turning response was calculated for a sliding window of 5 camera frames across all trials of a condition. The responses were averaged on a per-fly basis (see Figure 5 **b**), before being averaged across flies (top of Figure 5 **c,e,g**). Responses to counterclockwise stimulus movement were scaled by −1 and averaged together with the clockwise responses for the combined responses (Figure 5 **c, e**, bottom). The mean turning velocity during stimulus presentation is plotted as mean ± SEM across flies for the summary tuning curves (Figure 5 **d, f**).

## 3 RESULTS

### 3.1. Characterizing the technical performance of the experimental setup

As the inexpensive treadmill setup uses several components not typically used in animal behavior experiments, we measured many aspects of the system’s performance, and summarize thee results in Figure 4. To validate the Tablet’s display of our moving visual stimuli, we measured local brightness changes on one side of the display (position indicated on the right side of Figure 3 **b**) with a mounted photodiode (INL-3APD80, Inolux Corporation, Santa Clara, CA, USA). The data was viewed and logged on an oscilloscope (MDO3040, Tektronix Inc, Beaverton, OR, USA). Figure 4a, b shows typical measurements of the brightness changes measured for the moving patterns of the temporal frequency tuning conditions, and Figure 4 **c** shows typical measurements for moving patterns during the spatial frequency tuning conditions. These measurements suggest that the Fire 7 tablet reliably displays these periodic patterns, for example showing the expected periodic changes at the indicated temporal frequency. We note that even at the 30 Hz condition, which is half of the display refresh rate (lower trace in Figure 3 **b**), the stimulus timing looks extremely reliable; this condition is included as a stimulus control, since at half of the display refresh rate, the display flickers, and thus produces no net motion. The spatial frequency conditions show a similarly reliable periodic pattern at 7.5Hz (Figure 3**c**). The reduction in the sharpness of the edge transitions for smaller bars is simply due to spatial averaging by the sensor (and is a reasonable model for why the fly visual system also sees higher spatial frequency patterns as consisting of lower contrast levels).

During experiments, the FlyFlix client records the rendering status for each frame. Before a frame is displayed, the software asynchronously requests an update of the rendered content based on the current set of parameters. If this requests times out before the frame is rendered, then the previous content is shown again. Figure 4 **d** shows the percentage of frames that are rendered correctly and within the allotted time, which is on average the inter-frame-interval of ~17nis. We plot the percentage of correct frames for 33 open-loop experiments (from which the data of Figure 5 **a–d** were collected) as well as from 9 closed loop experiments (for which the behavioral data are not shown). For both configurations, the average performance is quite reliable, with more than 99.9 % of the frames correctly rendered. Figure 4 **e** provides details of the ~0.1% of cases when frames were not rendered in time. We do not find any systematic errors. On average one out of every one thousand frames skips exactly one frame update. Higher numbers of skipped frames are extremely rare, and tend to come in clusters, mostly during conditions with the same animals. Since FlyFlix records these measurements, trials above certain relevant thresholds can be identified post-hoc and removed from analysis.

Since the FlyFlix server FlyFlix client communicate via a network, we characterized the latency of this asynchronous bidirectional communication by sending timestamped packages from the server to the client, which immediately returns the package. In Figure 4 **f** we plot this WebSocket latency (WS) and also a ping using a lower network level (ICMP). 97.9% of the frames completed a round trip within 1 inter-frame-interval (~17 ms, indicated with vertical magenta lines in the plot) of the display, even though WebSocket based communication takes slightly longer with the additional protocol overhead. We expect that in our real application, the network reliability is even higher, since only half of a roundtrip is required to update the display.

In our experiments, we used our institute’s infrastructure: the FlyFlix server was connected to a wired network, the tablet connected via WiFi to a different subnet. Should the latency of an available network become too high, a local network router directly connecting FlyFlix server and client will improve the timing of the communication.

Taken together, the results of Figure 4, demonstrate that a low cost tablet provides a reliable visual display producing excellent stimulus control and timing over measured system events. These technical measurements show that our low cost system replaces many components typically required for precise experiment control (like Data Acquisition devices or high-end PC graphics cards) without sacrificing any performance, for the range of pattern speeds and network latencies described here.

### 3.2. Visually guided turning behaviors measured with the optimized setup

An important demonstration of our new, integrated system, is that ‘typical’ fly behaviors can be measured from flies tethered using our new tethering station and behavioral data collected using the new experimental setup. We focused on the optomotor responses, and present results from (33+20 =) 53 flies across 2 different protocols (detailed in sections 2.5 and 2.4). Figure 5 **a** shows the forward walking speed of flies during each trial of the temporal frequency protocol. Across speeds, flies walk with a similar speed, with a mean around 10 mm s^-1^ which is slightly higher than walking speeds measured in other fly-on-ball experiments (Creamer et al., 2018), and is only slightly slower than the walking speeds of freely walking flies at similar temperature (Ofstad et al., 2011).

When presented with rotating patterns, flies tend to turn in the direction of the pattern movement, a response we see in single trials and across trials for the example condition shown in 5 **b**. While there is some trial to trial variability, in nearly every trial, the flies turn in the clockwise, or positive direction (in tan) for clockwise pattern motion and in the counterclockwise, or negative direction (in purple) for counterclockwise pattern motion, a pattern that is clearly seen across flies and stimulus speeds (top of 5 **c**). The amplitude of the turning velocity we measure depends on the temporal frequency of the pattern movement (observable in the data combined from both directions, in the lower row of Figure 5 **c**). This is precisely the expected result, since temporal frequency tuning is a well described aspect of fly motion vision—insects are most sensitive to movement of periodic pattern with some temporal frequency optimum, and are less sensitive to movements with both higher and lower temporal frequency (Götz and Wenking, 1973). We compare our results, plotted using the mean responses during the period of stimulus presentation as a tuning curve, to the most relevant, recent independent measurement from another lab using a different setup (Figure 5 **d** contains an overlay of data from Creamer et al. (2018)). We find that in our experiments, for most conditions, flies are tuning more overall, and we see similar, monotonically increasing response levels up to 7.5 Hz motion. At the highest temporal frequencies we see an interesting difference– our responses fall off, while the responses from Creamer et al. (2018) remain much larger. We attribute this difference to limitations of our display, which as we have discussed, cannot properly display 30 Hz temporal frequency motion, and renders it as flicker. Overall, we find excellent concordance between our measurements and those of previous experimenters.

The optomotor turning response is also expected to depend on the spatial frequency of the grating pattern (Buchner, 1976; Creamer et al., 2018). We presented a series of grating patterns with different spatial periods at a fixed temporal frequency of 7.5 Hz. We observe large, consistent turning to patterns with a grating period above *λ* = 20° (Figure 5**e, f**). For narrower stripes, the responses are reduced, and in fact no consistent turning was measured for the pattern with *λ* = 5°. This result is expected based on prior work, and is remarkably similar to the measurement of Buchner (1976), who used a very different stimulus strategy.

Finally we tested the flies ability to track a moving bar, a behavior that is known to depend on both the motion and position of the moving object (Poggio and Reichardt, 1973; Bahl et al., 2013). As with the rotating grating patterns, we find that flies turn so as to follow the direction of the rotating bar (Figure 5 **g**). The peak turning velocity is similar between different rotational velocities of the stimulus, and quite similar to peak turning during the grating motion. To casually explore the position-dependence of the turning response, it suffices to note that most of the turning reaction occurs once the object (position indicated by the diagonal lines) has crossed the midline (most notable for *ω* = 90 ° s^-1^, 180 ° s^-1^, and 360 ° s^-1^. It is as if the flies don’t bother to orient towards an object they are likely to intercept as it approaches their midline, but once an object is getting away (as measured by its progressive, or front-to-back motion), the attempted tracking behavior rapidly increases. This response profile matches the recent measurements of walking flies (Bahl et al., 2013), but differs somewhat from the behavioral reactions of tethered flying flies that respond to both the regressive and progressive motion of the object (Reiser and Dickinson, 2010). For the fastest speeds tested, the flies cannot track, that is ‘catch up to’ the spinning bar, and the responses are seen to lag the position of the stripe by more than 100 ms. The flies’ turning velocity almost disappears at the condition with the stimulus angular velocity of *ω* = 1350 ° s^-1^. In this condition, the object moves across the 60 fps display in less than 8 frames with displacements of more than 20° between frames, which are too large for the fly to smoothly integrate as motion. Consequently, the turning response is barely noticeable.

In Figure 5 we summarize the behavior of *Drosophila* in our optimized, inexpensive treadmill setup, in a sophisticated range of stimulus conditions. We show clear symmetric turning responses to all symmetric stimulus conditions. The temporal and spatial frequency tuning as well as the object tracking behaviors are highly similar to previously published measurements from other labs using different experimental setups. Based on these results, we unreservedly recommend this low-cost setup, not only for teaching purposes, but for nearly any research application.

## 4 DISCUSSION

In this paper we have described our re-implementation of a complete system for tethering flies and the accompanying experimental setup for measuring tethered fly walking behavior to controlled visual stimuli (Figures 1,2). Our spherical treadmill setup takes a fresh look at the fly-on-a-ball paradigm. While the design is guided by several decades of experimental methods development, we are intensely focused on optimizing the setup by simplifying the components, reducing costs, and ensuring availability. Since many of the components have not previously been deployed in animal behavior setups, we validated the performance of the display (Figure 4). We found excellent reliably for the low-cost display and low network latencies, which combine to establish a highly reliable new method for experimental control. This system comes with other advantages such as a flexible stimulus control software that can dynamically correct for the viewing angle (Figure 3). Finally, we measured the walking behavior of flies to a range of moving visual stimuli and confirmed, in exquisite detail, that our new setup is capable or reproducing nearly all relevant prior measurements using similar visual stimuli. Based on this experience we believe that our setup will be ideal for use in teaching courses and for a wide range of laboratory uses. We sincerely hope that the reduced complexity and enhanced accessibility of these setups will excite many young scientists about quantitative animal behavior, and will increase the reproducibility of research observations. In the following sections we discuss cost savings of our system, the cost of cost savings in the form of limitations, some possible extensions and future work.

### 4.1. Costs and Availability

We cannot claim a system to be inexpensive without estimating the cost of acquiring such a system, and also comparing it to a contemporary alternative. We have estimated the cost based on building a single setup, using parts available in the US, during the spring of 2021. Many of the components are available as generic parts from multiple vendors, we have selected example sources to illustrate the price range for potential cost savings and overall costs. We provide links for the same purpose, and these are not an endorsements for or against particular vendors, especially since the products might not be available in different countries. We estimate the prices for 3D printed components using the online instant quote at https://craftcloud3d.com. For the laser cutting, we use estimates from https://ponoko.com. Those with access to a 3D printer or a laser cutter can expect to save a little more money.

For the comparison to a contemporary setup, we surveyed several groups and specified a system that would realistically represent the type of setup we would build in our lab today for ongoing research projects. Below we detail a few key components, and summarize the systems’ cost in Table 1 and Table 2, and in Figure 6). Figure 6 shows that we can assemble both complete systems for ~$300. whereas the standard, yet very nice, pair of setups would cost ~$15,000, a remarkable 50-fold cost reduction.

**Figure 6.**
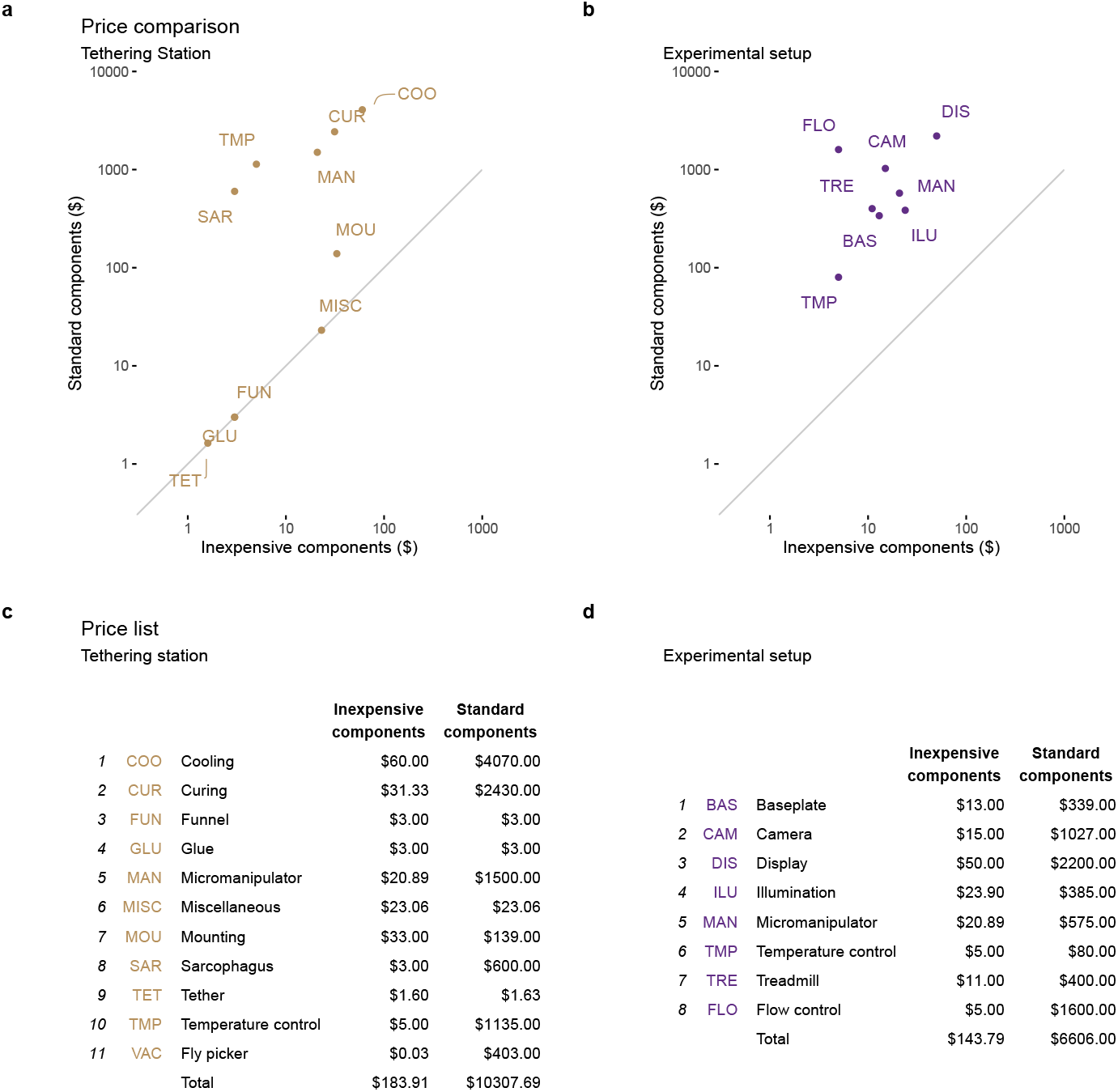
Estimated cost savings for each setup. The price of each functional unit is compared between the standard setups and the optimized, inexpensive tethering station **a** and experimental (treadmill) setup **b**. The line in each of **a,b** represents an equal price in both setups. In **c** and **d** we list the components (or functional units) represented by the labels in **a** and **b**. We estimate a roughly 50-fold cost savings between the two versions of these systems. Further details are provided in the text and in the accompanying tables of components

**Table 1.**
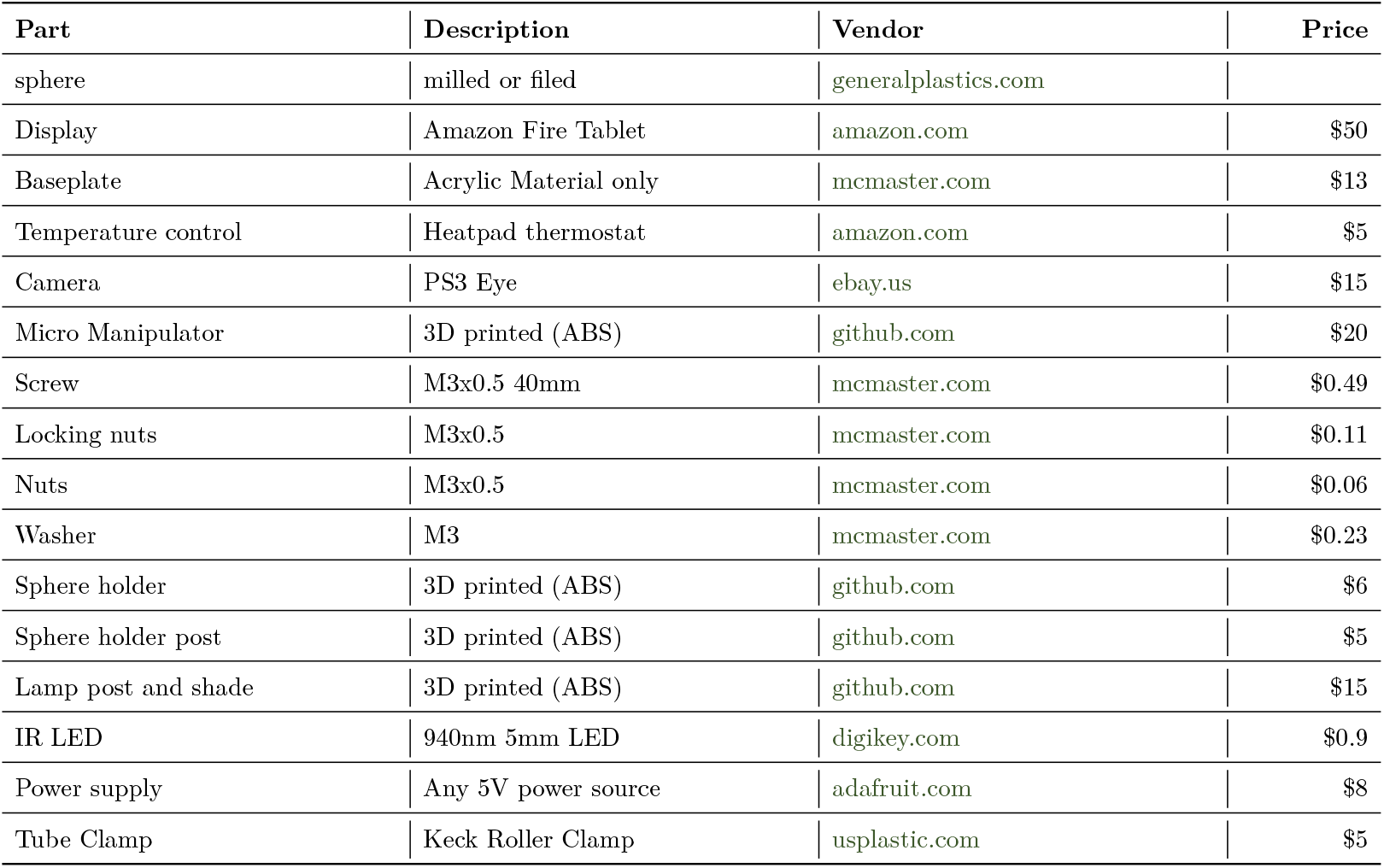
Price estimation of parts for Experimental setup.

**Table 2.** Suggestion for components in an inexpensive treadmill tethering station.

| Part | Description | Vendor | Price |
| --- | --- | --- | --- |
| microscope | various; any dissecting scope |  |  |
| glue | glass to glass adhesive | kemxert.com | \$3 |
| UV protective glasses | many | | \$30 |

| Part | Description | Vendor | Price |
| --- | --- | --- | --- |
| Fly picker wand | Transfer Pipette | amazon.com | \$0.03 |
| Temperature control | Chiller peltier thermostat | amazon.com | \$5 |
| Tether | Blunt tip dispensing needle 34G | bstean.com | \$1.6 |
| Funnel | 3D printed (PLA, ABS) | github.com | \$3 |
| Micro Manipulator | 3D printed (ABS) | github.com | \$20 |
| Screw | M3x0.5 40mm | mcmaster.com | \$0.49 |
| Locking nuts | M3x0.5 | mcmaster.com | \$0.11 |
| Nuts | M3x0.5 | mcmaster.com | \$0.06 |
| Washer | M3 | mcmaster.com | \$0.23 |
| Heat pump | Peltier element on heat sink | adafruit.com | \$35 |
| Heat pump power supply | 12V 5A | adafruit.com | \$25 |
| Sarcophagus | 3D printed (ABS) | github.com | \$3 |
| Tether Station holder | Laser cut (acrylic) | github.com | \$15 |
| UV Curing light | UV Keychain light | amazon.com | \$1.33 |
| Paintbrush | NA | amazon.com | \$7 |
| Round bottom tube | Chilling tube | mcmaster.com | \$0.06 |
| Heat sink holder | Hand rest | ponoko.com | \$18 |
| Hollow Body Pin Vise | Flyhook holder | mcmaster.com | \$16 |

Software: A simple way to reduce costs and increase access is to exclusively use open-source software. From GNU/Linux as the operating system, to FicTrac, camera drivers, FlyFlix, and Firefox, all are available without paying software licenses. Furthermore, the majority of the components in the Component-Designs GitHub repository are constructed using freely available software such as FreeCAD, KiCAD, and Inkscape, which enables the unrestricted exchange of design files.

Baseplate: We used an acrylic board with a grid of holes, which can save more than $100 per setup compared to a professional optomechanical board. If available, a laser cutter is ideal for fabricating these boards within minutes. In the accompanying repository we provide files for laser cutter, CNC-machines, or as a blueprint for hand-drilling.

Camera: We use a PlayStation Eye as the input for FicTrac. We could not find another widely available low cost camera that achieves high frame rate imaging. Since it is mass-produced as a toy, they are available at different vendors and secondary markets for around $5 to $20. We describe the required modifications with a macro lens ($15) and M12 lens holder ($2) in the supplements. This modification takes between 30 min and 60 min (see Figures S1 and S2).

Micromanipulator: Finding low-cost alternatives for this enabling tool was one of the initial drivers of this project. With the purchase of three-axis linear stages at an overseas vendors, cost savings of up to 80 % on that one part enabled us to build several setups for the teaching course (shown in Figure S3). Nevertheless, at $80 to $100, they are still a considerable expense, especially if using two per setup. For labs with access to a 3D printer, our design of the micromanipulator costs less than $5 in materials (including nuts and screws). Ordering the parts through 3D printing services increases the cost to around $15. In addition to printing time, ~ 1 h of build time is required. Similarly the Sphere holder is easy to adapt and cheap to print with material costs below $3 and online services charging around $6.

### 4.2. Trade-offs and limitations

The flexibility and modularity of our proposed system is also a limitation: it takes more time and effort to make and assemble the systems based on components from multiple vendors, rather than ordering readymade products. We sought to replace all custom parts with commercially available inexpensive components wherever possible, such as the display system or the tethers, but in many cases, no alternative existed and we turned to custom designs.

Many components of our setup are produced in a 3D printer or a laser cutter. This may increase access compared to custom-machines metal parts, but it is still a limitation. Nevertheless, we see three main alternatives to access these parts: (1) high quality 3D printers are becoming more affordable and easier to use, (2) maker spaces provide access to 3D printers in communities across the world, and (3) many companies offer 3D prints as a service. We used the third (and most expensive) option in our cost estimates (Figure 6). We consider access to a laser cutter as nice, but not necessary for building this setup (alternatives discussed throughout). The factors regarding price, maker spaces, and online services also apply to laser-cutting acrylics. Building a new experimental setup is always a time-consuming endeavor, but even more so when the components need to be built from scratch. We estimate ~5 h of printing time on the Stratasys F-170 printer. We further estimate that another ~5h are necessary for assembling the first setup. Times will vary greatly and assembly will speed up over time.

FlyFlix, the system of a single server providing stimuli for network connected display clients, is extensible to many displays. Nevertheless, for the current description we limited the use to a single display in front of the fly covering ~130° in azimuth and 100° in elevation. In our standard fly behavioral setups, we prefer to stimulate larger portions of the fly’s field of view, especially laterally. Furthermore, the tablet we chose only supports refresh rates of 60 fps. This limits the speed of stimuli that can be shown, including to motion speeds that the fly can perceive (see Figure 5 **d**). Newer handheld displays with higher refresh rates and gaming monitors used in other experimental setups overcome this limitation, but at significantly increased cost (Kaushik et al., 2020). The FlyFlix software is agnostic to the display and should work out of the box with higher refresh rates. Nevertheless, network latency will be a limiting factor for high-speed closed-loop systems, but there is little reason to belive that flies (or just about any animal) required closed loop latencies that are less than ~10ms.

### 4.3. Extensions and future work

The inexpensive treadmill project was inspired by the challenge of setting up multiple rigs in a teaching lab to provide hands-on experience with *Drosophila’s* fascinating walking responses to visual stimuli. The setup has been optimized and so far only tested with fruit flies. Nevertheless, we expect that adapting the setup to other insects should be straightforward. The Sarcophagus already accommodates many body sizes and could be modified for others. A much larger insect may require a larger ball size, but fortunately, the nature of our manufacturing process and the availability of our 3D designs allows any components to be scaled to adapt to specific animal sizes.

While we have focused on visually guided behaviors with this setup, it would be very exciting to implement other types of sensory stimulation: wind, humidified air, sounds, odors, or even polarized light (Mathejczyk and Wernet, 2020). All of these can be integrated into our experimental design with little to no modification to the existing components.

While we have achieved all of our initial goals, we continue working to improve the system. We are testing more accessible alternatives to our hand-filed balls and seeking a good replacement for laboratory wall air to float the ball. While we have implemented closed-loop protocols and confirmed that they are technically working, we have so far not been impressed with the behavioral results from these trials, and so we continue to optimize those methods. Finally we will implement a low-cost solution for optogenetic stimulation of the walking flies, and will adapt the setup as needed so that it can be mounted under a microscope to accommodate electrophysiology or imaging. All updates will be posted on the accompanying repository and we welcome feedback, ideas, and contributions from the research community.

## ACKNOWLEDGMENTS

We wish to thank the *Drosophila* Neurobiology: Genes, Circuits & Behavior course (and organizers E. Heckscher, A. Keene, and A. Frank) for inspiring us to work on this project. We thank R. Franconville for using and providing feedback on an early iteration of the setup in a summer course. H. Haberkern for FicTrac consultation, I. Negrashov and J. Talbot for engineering assistance, E. Gruntman for advice and testing, and E. Rogers for help with flies. We are also grateful to members of the Reiser Lab for comments on the work and manuscript.

## AUTHOR CONTRIBUTIONS

M.B.R. and F.L. conceived of the project and wrote the paper together. F.L. developed methodology, wrote software, carried out experiments, and analyzed data.

## Supplementary Material for

**Figure S1.**
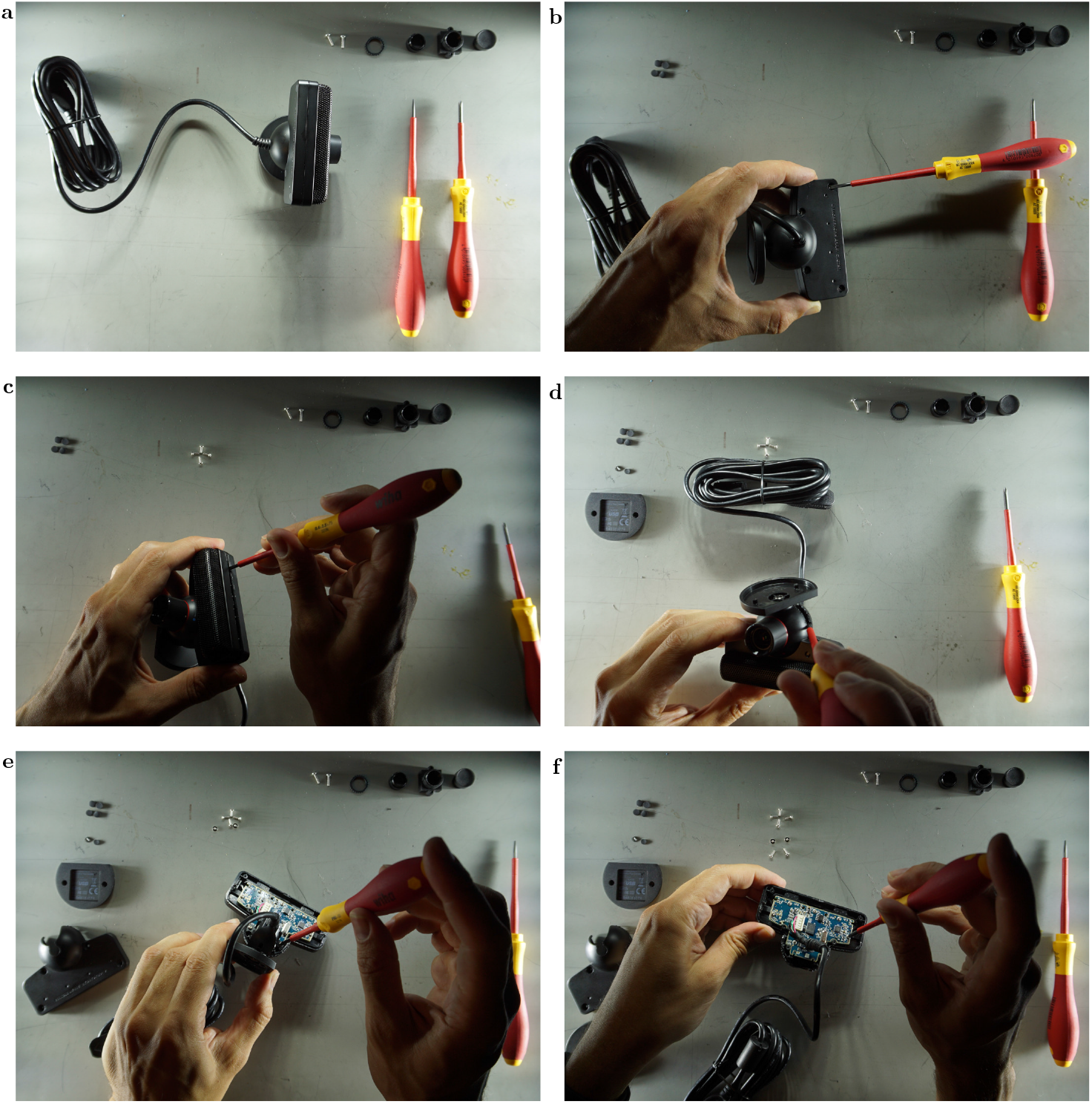
Disassembling the camera: **a**: Tools and parts required for modification. **b**: Unscrew the back of the camera. **c, d**: Carefully break notches holding back and front together. **e, f**: Unscrew cable holder and PCB.

**Figure S2.**
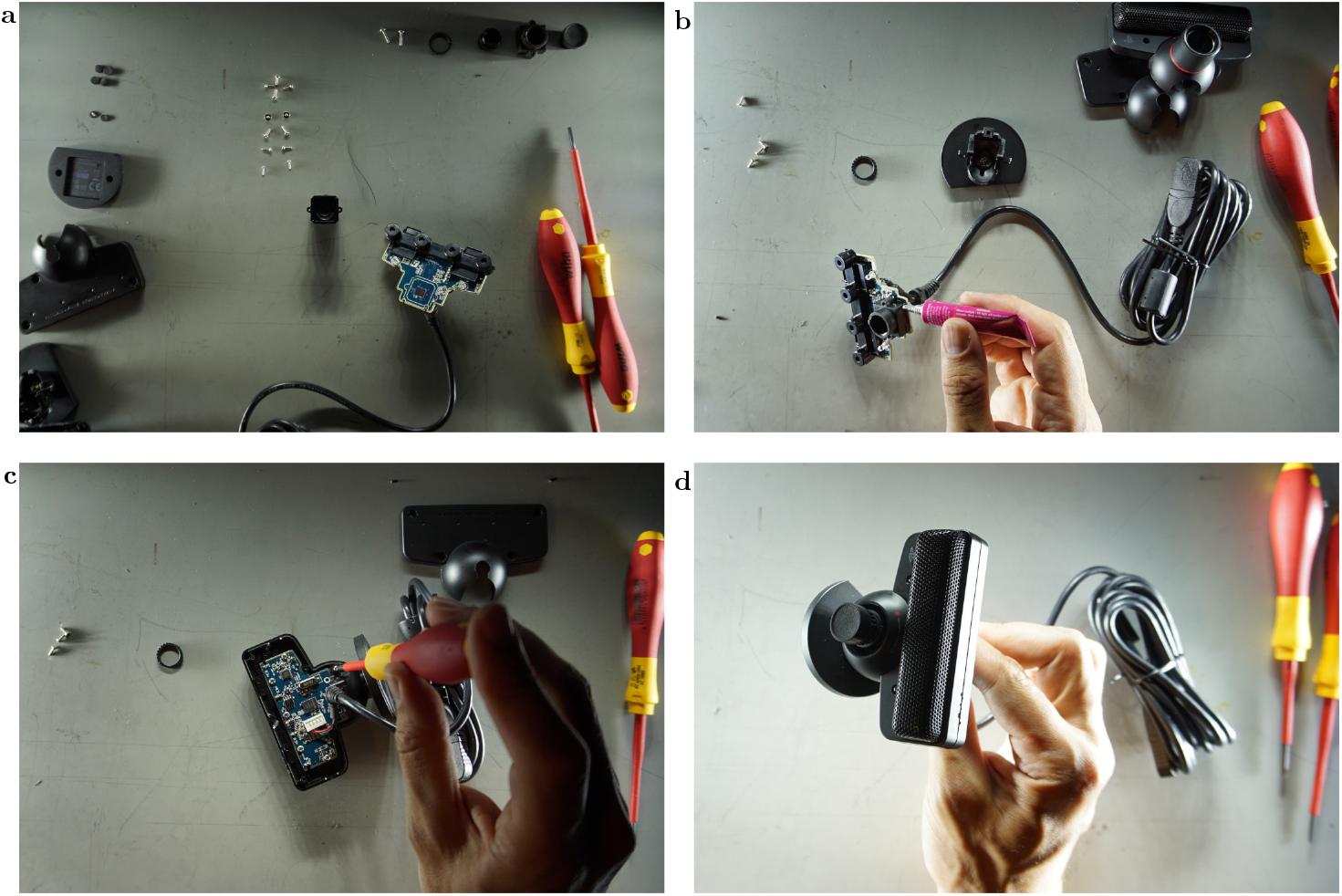
Modifying the camera: **a**: remove the lens from the front of the PCB. **b**: Attach the M12 lens mount in the same place. **c**: reattach PCB to camera case. **d**: Close camera case. Without the broken notches removed in Figure S1 c, some glue is needed. In the picture, the macro lens has a lens cover.

**Figure S3.**
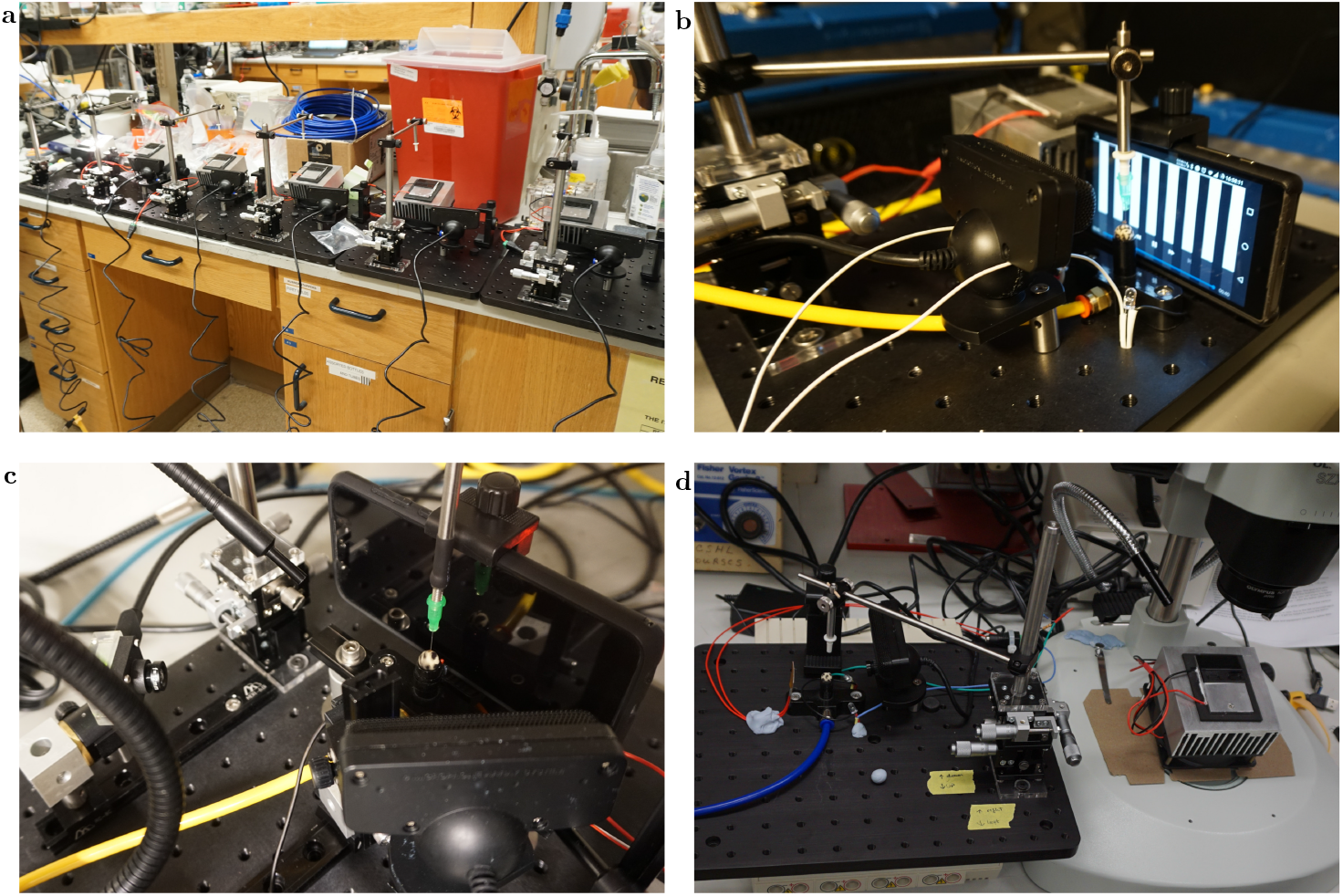
Early versions of these setups: **a**: 6 setups used in the CSHL course. **b**: Shared Micromanipulator in combined tethering and experimental setup. **c** Prototyping camera angles and display sizes **d**: Shared Micromanipulator between tethering and experiment, for a heating setup, without the display.

**Figure S4.**
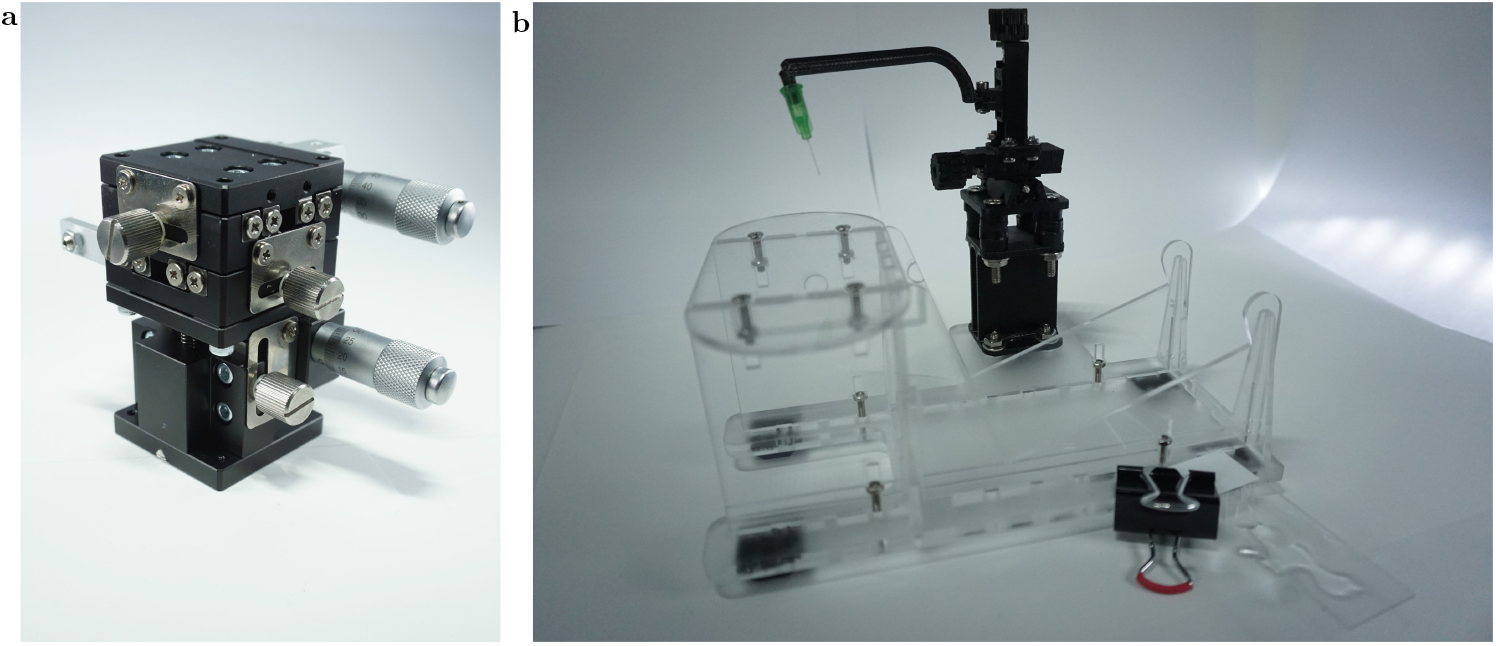
Micromanipulators: **a**: LD40-LM micromanipulator from overseas distributor. **b**: Arm rest with fixture to hold the Chiller at the ideal angle, cut from acrylic. A glass slide for holding a few drops of glue is in the foreground and the 3D-printed micromanipulator in the background.

**Figure S5.**
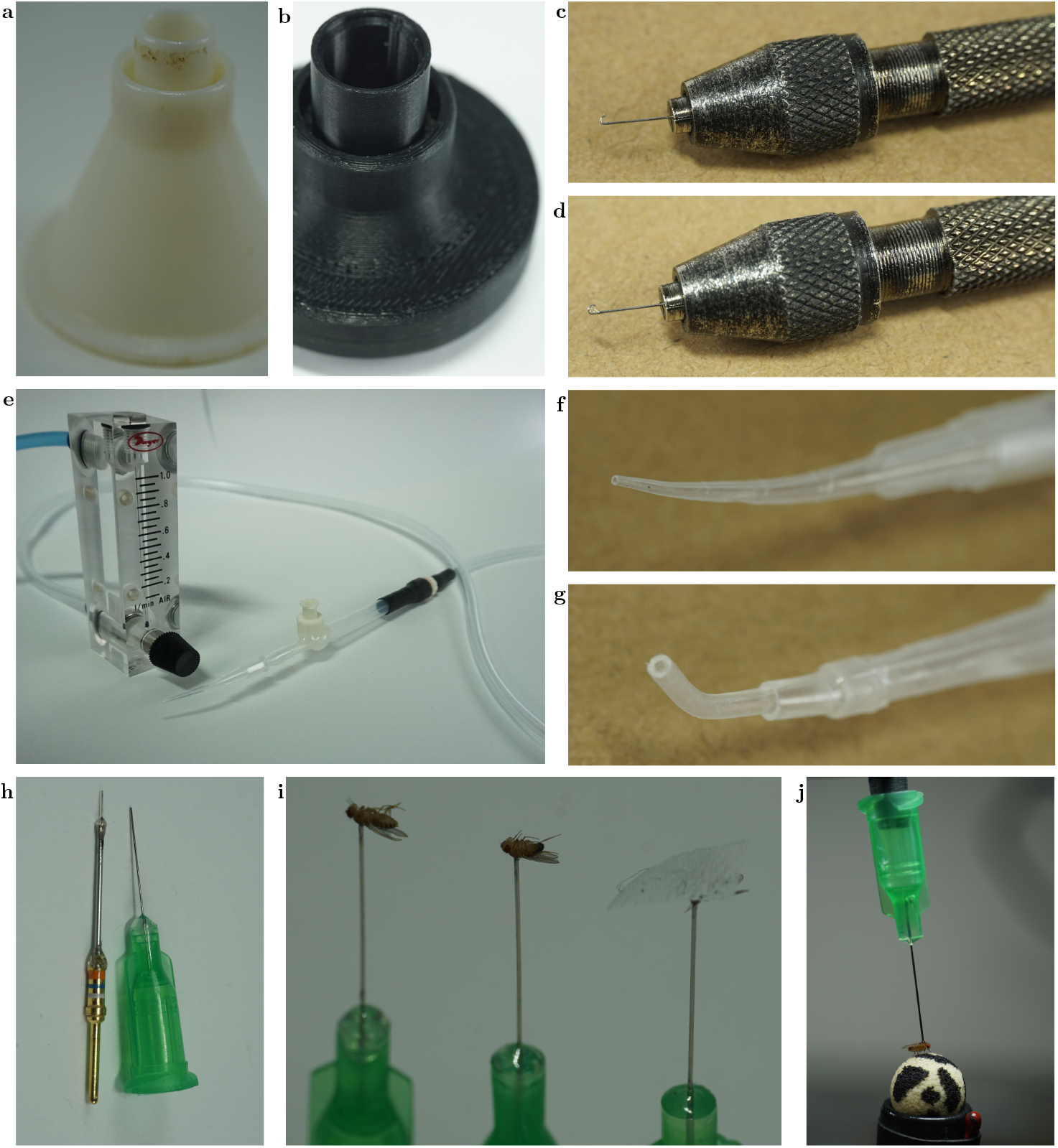
Gallery of useful tools: **a, b**: 3D-printed funnels used to move flies between bottles and vials. A bent minutien pin used to move the fly in the Sarcophagus and apply glue, without glue **c** and **d** with glue. **e** Flowmeter to regulate negative pressure for the fly picker. Tip of the fly picker made from micropipette tip (**f**) and shrink tube (**g**). **h** Traditionally made tether (left) and blunt dispensing tip with Luer lock alternative (right). **i** tethered flies in holding area prior to experiments, the 3rd holding a piece of tissue. **j** Body-fixed, tethered fly on a ball, showing the modest ‘uphill’ orientation of the fly

## TABLES

**Table S1.** Components used in the tethering station for the inexpensive treadmill. The exact links and prices may vary, but with the product number (PN) and description, other vendors and similar products can be found.

| part | description | link | PN | price |
| --- | --- | --- | --- | --- |
| microscope | various; any dissecting scope |  |  |  |
| glue | glass to glass adhesive | http://www.kemxert.com/product-catalog.cfm?pg=product-catalog&prd_pct_id=2 | KOA-300 | \$3 |
| UV protective glasses | many | | | \$30 |
| Fly picker wand | Transfer Pipette | https://amazon.com/dp/B07RFFMKBM | | \$0.03 |
| Temperature control | Chiller peltier thermostat | https://amazon.com/dp/B07VDRGK9F | XH-W1209 | \$5 |
| Tether | Blunt tip dispensing needle 34G | https://www.bstean.com/product/industrial-unsterilized-blunt-tip-dispensing-needle-with-luer-lock-34-ga-x-1-2-50-pcs/ | AG-ABSS-99D0 | \$1.6 |
| Funnel | 3D printed (PLA, ABS) | https://reiserlab.github.io/Component-Designs/tether/funnels | | \$3 |
| Micro Manipulator | 3D printed (ABS) | https://reiserlab.github.io/Component-Designs/tether/micromanipulator | | \$20 |
| Screw | M3x0.5 40mm | https://www.mcmaster.com/91287A028/ | 91287A028 | \$0.49 |
| Locking nuts | M3x0.5 | https://www.mcmaster.com/90576A102/ | 90576A102 | \$0.11 |
| Nuts | M3x0.5 | https://www.mcmaster.com/90592A085/ | 90592A085 | \$0.06 |
| Washer | M3 | https://www.mcmaster.com/95610A530/ | 95610A530 | \$0.23 |
| Heat pump | Peltier element on heat sink | https://www.adafruit.com/product/1335 | 1335 | \$35 |

| part | description | link | PN | price |
| --- | --- | --- | --- | --- |
| Heat pump power supply | 12V 5A | https://www.adafruit.com/product/352 | 352 | \$25 |
| Sarcophagus | 3D printed (ABS) | https://reiserlab.github.io/Component-Designs/tether/sarcophagus | | \$3 |
| Tether Station holder | Laser cut (acrylic) | https://reiserlab.github.io/Component-Designs/tether/station | | \$15 |
| UV Curing light | UV Keychain light | https://amazon.com/dp/B086PW9DBH | | \$1.33 |
| Paintbrush | NA | https://amazon.com/dp/B0878MN2VR | | \$7 |
| Round bottom tube | Chilling tube | https://www.mcmaster.com/7012A54/ | 7012A54 | \$0.06 |
| Heat sink holder | Hand rest | https://www.ponoko.com/materials/anti-static-acrylic | | \$18 |
| Hollow Body Pin Vise | Flyhook holder | https://www.mcmaster.com/8455A18/ | 8455A18 | \$16 |

**Table S2.** Components used in the inexpensive treadmill experimental setup. The exact links and prices may vary, but with the product number (PN) and description, other vendors and similar products can be found. Some parts are listed as alternatives, for example the commercially available LD40-LM micromanipulator and the 3D-printed one.

| part | description | link | PN | price |
| --- | --- | --- | --- | --- |
| sphere | milled or filed | https://www.generalplastics.com/products/fr-7100 | FR-7110 |  |
| Display | Amazon Fire Tablet | https://amazon.com/dp/B07FKR6KXF | KFMUWI | \$50 |
| Baseplate | Acrylic Material only | https://www.mcmaster.com/4615T37 | 4615T37 | \$13 |
| Baseplate | 5.5mm acrylic with holes | https://www.ponoko.com/materials/black-acrylic | | \$27 |

Table S2 Components used in the inexpensive treadmill experimental setup.
| part | description | link | PN | price |
| --- | --- | --- | --- | --- |
| Micro Manipulator | XYZ micromanipulator | https://www.aliexpress.com/item/33013923564.html | LD40-LM | \$80 |
| Flow meter | Dwyer Flowmeter | https://www.dwyer-inst.com/Product/Flow/Flowmeters/VariableArea/SeriesVF | VFA-23 | \$40 |
| Temperature control | Heatpad thermostat | https://amazon.com/dp/B07VDRGK9F | XH-W1209 | \$5 |
| Camera | PS3 Eye | https://ebay.us/8Gwt0X | 7010571 | \$15 |
| Micro Manipulator | 3D printed (ABS) | https://reiserlab.github.io/Component-Designs/tether/micromanipulator | | \$20 |
| Screw | M3x0.5 40mm | https://www.mcmaster.com/91287A028/ | 91287A028 | \$0.49 |
| Locking nuts | M3x0.5 | https://www.mcmaster.com/90576A102/ | 90576A102 | \$0.11 |
| Nuts | M3x0.5 | https://www.mcmaster.com/90592A085/ | 90592A085 | \$0.06 |
| Washer | M3 | https://www.mcmaster.com/95610A530/ | 95610A530 | \$0.23 |
| Sphere holder | 3D printed (ABS) | https://github.com/reiserlab/Component-Design | | \$6 |
| Sphere holder post | 3D printed (ABS) | https://reiserlab.github.io/Component-Designs/walking/sphere-holder | | \$5 |
| Lamp post and shade | 3D printed (ABS) | https://reiserlab.github.io/Component-Designs/walking/illumination | | \$15 |

Table S2 Components used in the inexpensive treadmill experimental setup.
| part | description | link | PN | price |
| --- | --- | --- | --- | --- |
| IR LED | 940nm 5mm LED | https://www.digikey.com/short/5hnfw5bq | IR204 | \$0.9 |
| Power supply | Any 5V power source | https://www.adafruit.com/product/276 | 276 | \$8 |
| Tube Clamp | Keck Roller Clamp | https://www.usplastic.com/catalog/item.aspx?itemid=31260 | 16004 | \$5 |

**Table S3.**
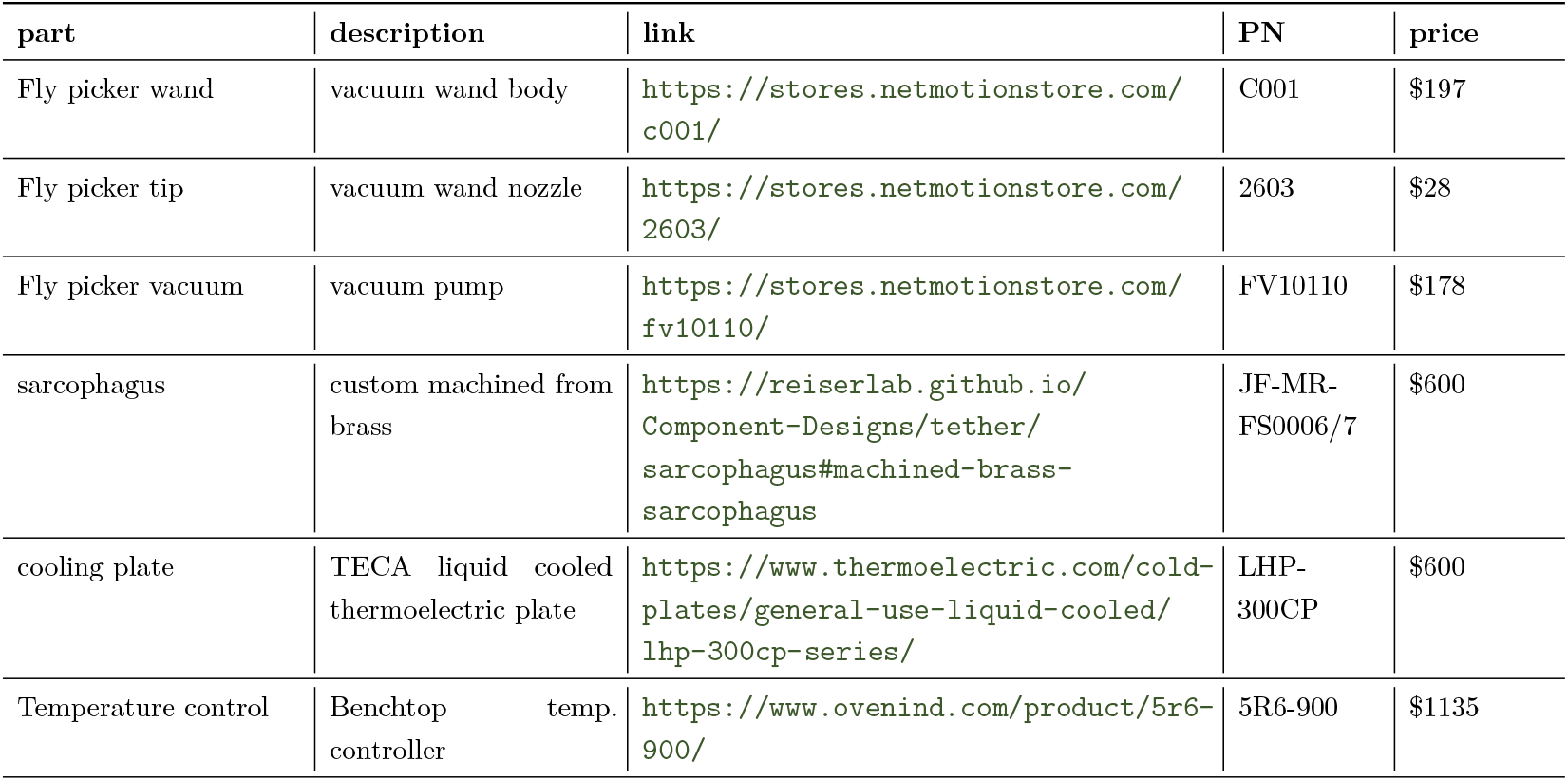
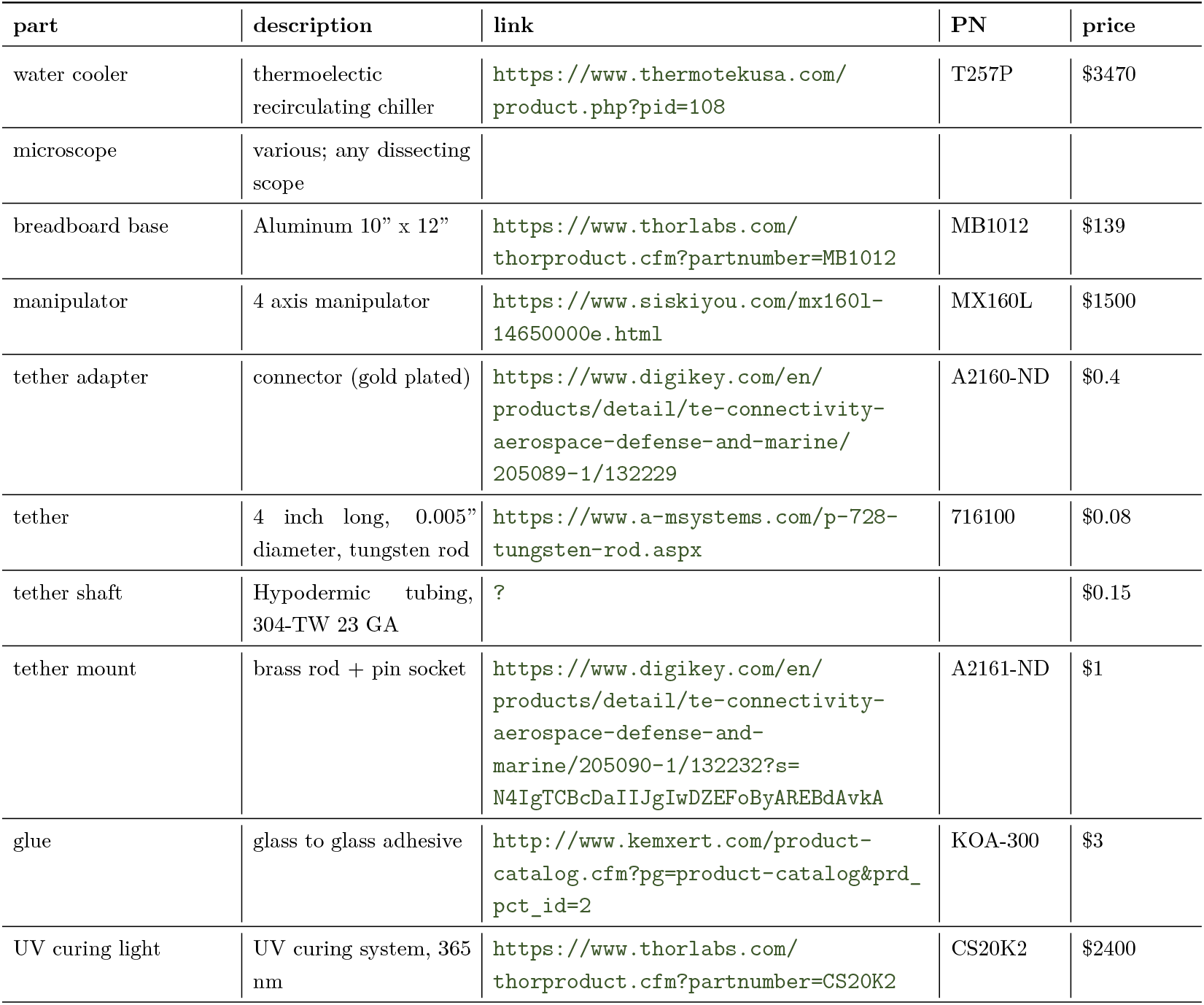
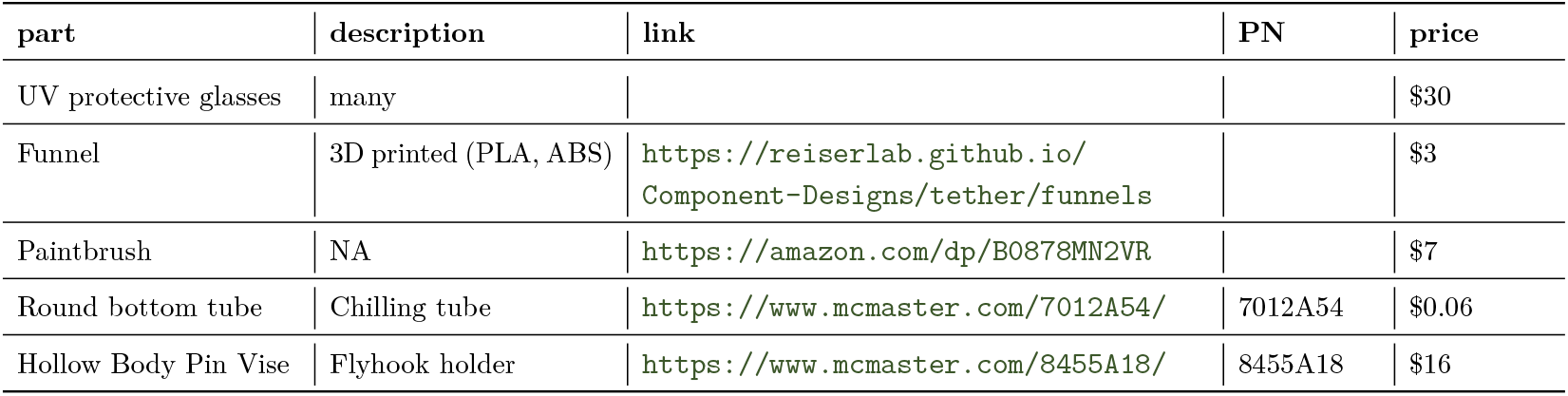
Examples for components used in typical preparatory setups.

**Table S4.**
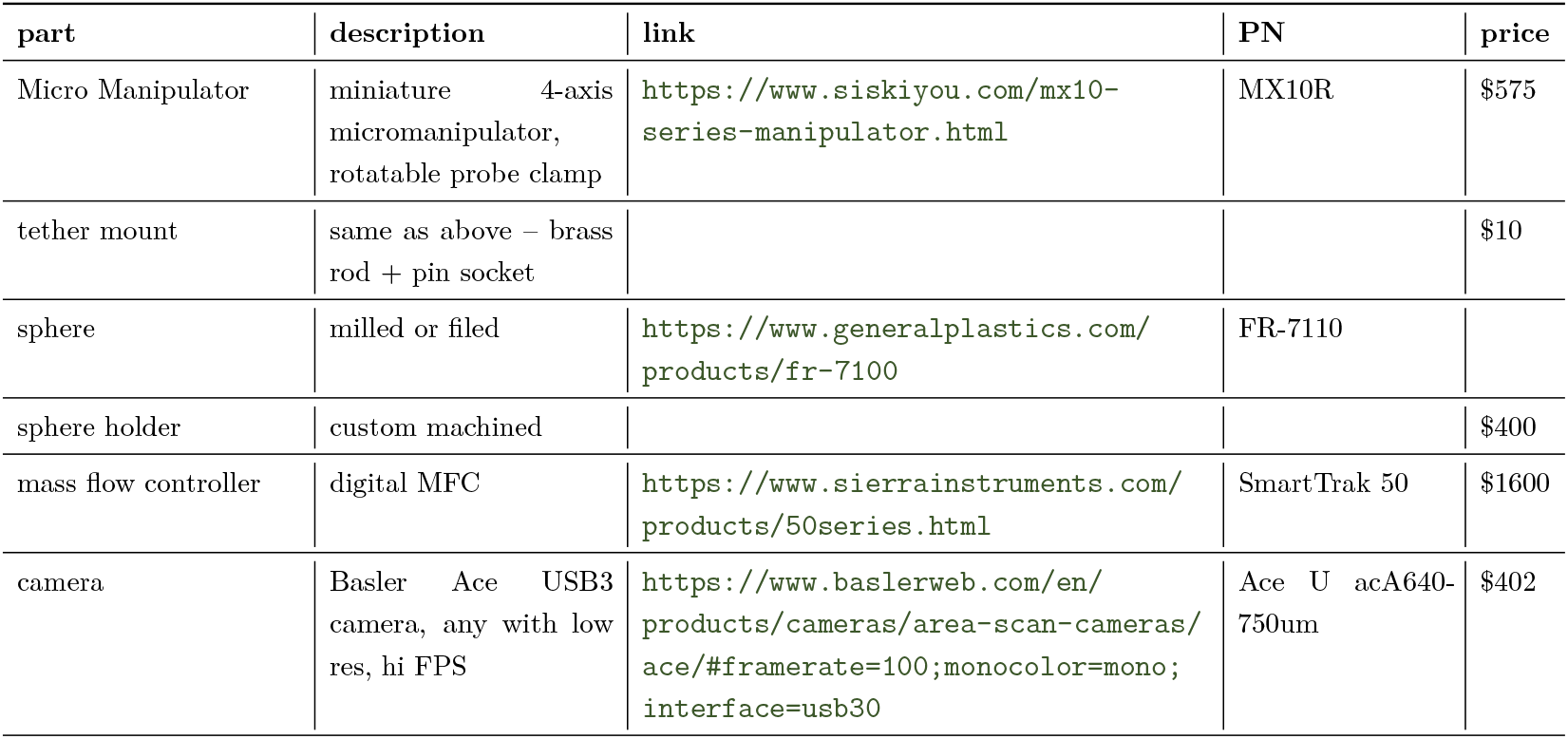
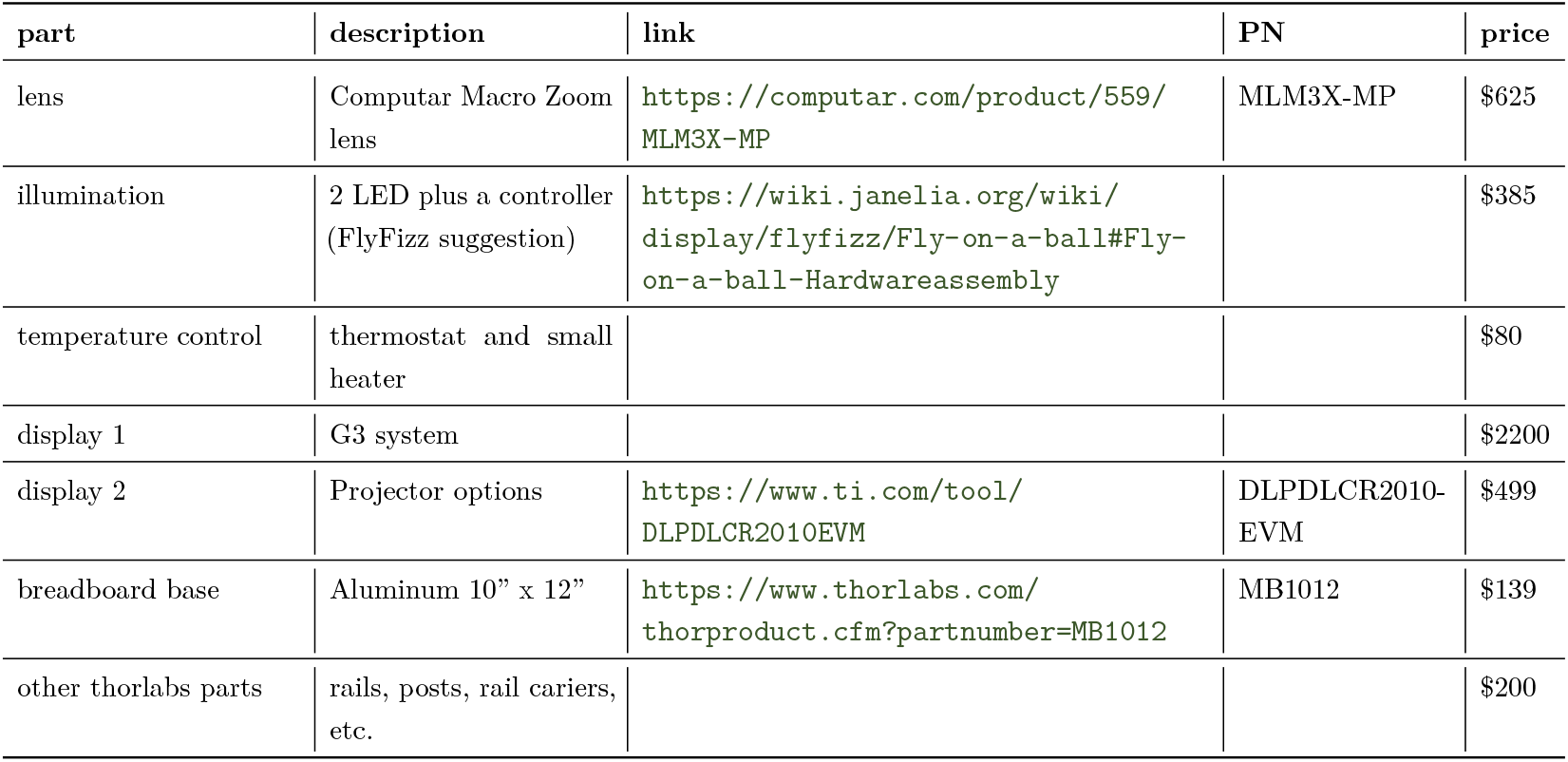
Examples for components used in typical experimental setups.

| part | description | link | PN | price |
| --- | --- | --- | --- | --- |
| Micro Manipulator | miniature 4-axis micromanipulator, rotatable probe clamp | https://www.siskiyou.com/mx10-series-manipulator.html | MX10R | \$575 |
| tether mount | same as above – brass rod + pin socket | | | \$10 |
| sphere | milled or filed | https://www.generalplastics.com/products/fr-7100 | FR-7110 |  |
| sphere holder | custom machined | | | \$400 |
| mass flow controller | digital MFC | https://www.sierrainstruments.com/products/50series.html | SmartTrak 50 | \$1600 |
| camera | Basler Ace USB3 camera, any with low res, hi FPS | https://www.baslerweb.com/en/products/cameras/area-scan-cameras/ace/#framerate=100;monocolor=mono;interface=usb30 | Ace U acA640-750um | \$402 |

Table S4 Examples for components used in typical experimental setups.
| part | description | link | PN | price |
| --- | --- | --- | --- | --- |
| lens | Computar Macro Zoom lens | https://computar.com/product/559/MLM3X-MP | MLM3X-MP | \$625 |
| illumination | 2 LED plus a controller (FlyFizz suggestion) | https://wiki.janelia.org/wiki/display/flyfizz/Fly-on-a-ball#Fly-on-a-ball-Hardwareassembly | | \$385 |
| temperature control | thermostat and small heater | | | \$80 |
| display 1 | G3 system | | | \$2200 |
| display 2 | Projector options | https://www.ti.com/tool/DLPDLCR2010EVM | DLPDLCR2010-EVM | \$499 |
| breadboard base | Aluminum 10" x 12" | https://www.thorlabs.com/thorproduct.cfm?partnumber=MB1012 | MB1012 | \$139 |
| other thorlabs parts | rails, posts, rail carriers, etc. | | | \$200 |

